# Full-Length Context Disrupts Folding of IgG-Binding Domains of Protein A

**DOI:** 10.64898/2025.12.08.692973

**Authors:** Kosar Rahimi, Albert Halbing, Minh Ngoc Nguyen, Mehmet Sen, Richard C. Willson, Gül H. Zerze

## Abstract

Multidomain proteins are often thought to fold as collections of independently stable domains, a modularity that underpins many assumptions in structural biology and design. Here, we challenge this view by examining the folding behavior of full-length Staphylococcal protein A (SpA), a 516-residue multidomain protein containing five immunoglobulin (Ig)-binding domains. Although each of the five Ig-binding domains of SpA folds stably in isolation (as it is already known experimentally and also confidently predicted by AI models), here, we show that the full-length construct and the individual Ig-binding domains in the full-length construct fail to adopt a stable three-dimensional structure in solution (despite being predicted to be folded by AI models). Instead, full-length SpA populates a compact yet predominantly disordered ensemble with residual secondary structure, where the folded state of each Ig-binding domain is thermodynamically unfavorable. These findings not only challenge long-held assumptions about the modular architecture and stability of SpA but also underscore the limitations of AI-based predictors when decoupled from the thermodynamic context. This work has implications for validating structure predictions, understanding multidomain architecture, and designing modular proteins for biotechnology and medicine.

## Introduction

Predicting how proteins gain function through structure remains a central goal of molecular biology, with wide-ranging implications for understanding cellular function, engineering new biomolecules, and treating disease.^1–4^ Deep learning tools such as AlphaFold have transformed our ability to predict folded protein structure from sequence, offering confident predictions even for large and complex systems. However, these predictions reflect fundamentally static pictures and remain context-agnostic: they do not account for solvent, temperature, pH, folding equilibria, or the dynamic ensemble of conformations that proteins adopt in solution.^5–7^ The underlying thermodynamic landscape must be resolved to understand how proteins behave in solution. This is particularly crucial for protein design and therapeutic development, where structural stability, disorder, and dynamics determine biological outcome.^8,9^

Multidomain proteins pose a particular challenge. Many studies have characterized the folding of isolated domains or fragments,^10–13^ but most proteins are multidomain in nature. Multidomain proteins may exhibit emergent properties due to long-range coupling and are not readily decomposable into modular parts.^3,14–19^ Although individual domains often fold stably in isolation, interdomain contacts, linker flexibility, and crowding effects can shape their behavior in the context of a full-length protein. Capturing this behavior requires going beyond static structures toward thermodynamically-informed ensemble descriptions.

Among multidomain proteins, Staphylococcal Protein A (SpA) from *Staphylococcus aureus* (*S. aureus*) exemplifies this modular paradigm. This bacterial surface protein consists of five tandem homologous immunoglobulin (Ig)-binding domains (E, D, A, B, and C), along with a C-terminal cell wall domain (domain X) (Fig. 1a). It acts as a critical virulence factor by interacting with multiple host targets: it binds the Fc region of IgG to block opsonization,^20^ engages VH3-type B cell receptors to induce apoptosis,^21,22^ activates TNFR1-driven inflammation,^23^ and promotes adhesion via von Willebrand factor.^24^ These multifaceted roles underscore SpA’s importance in immune evasion and pathogenesis.^25,26^

**Fig. 1.**
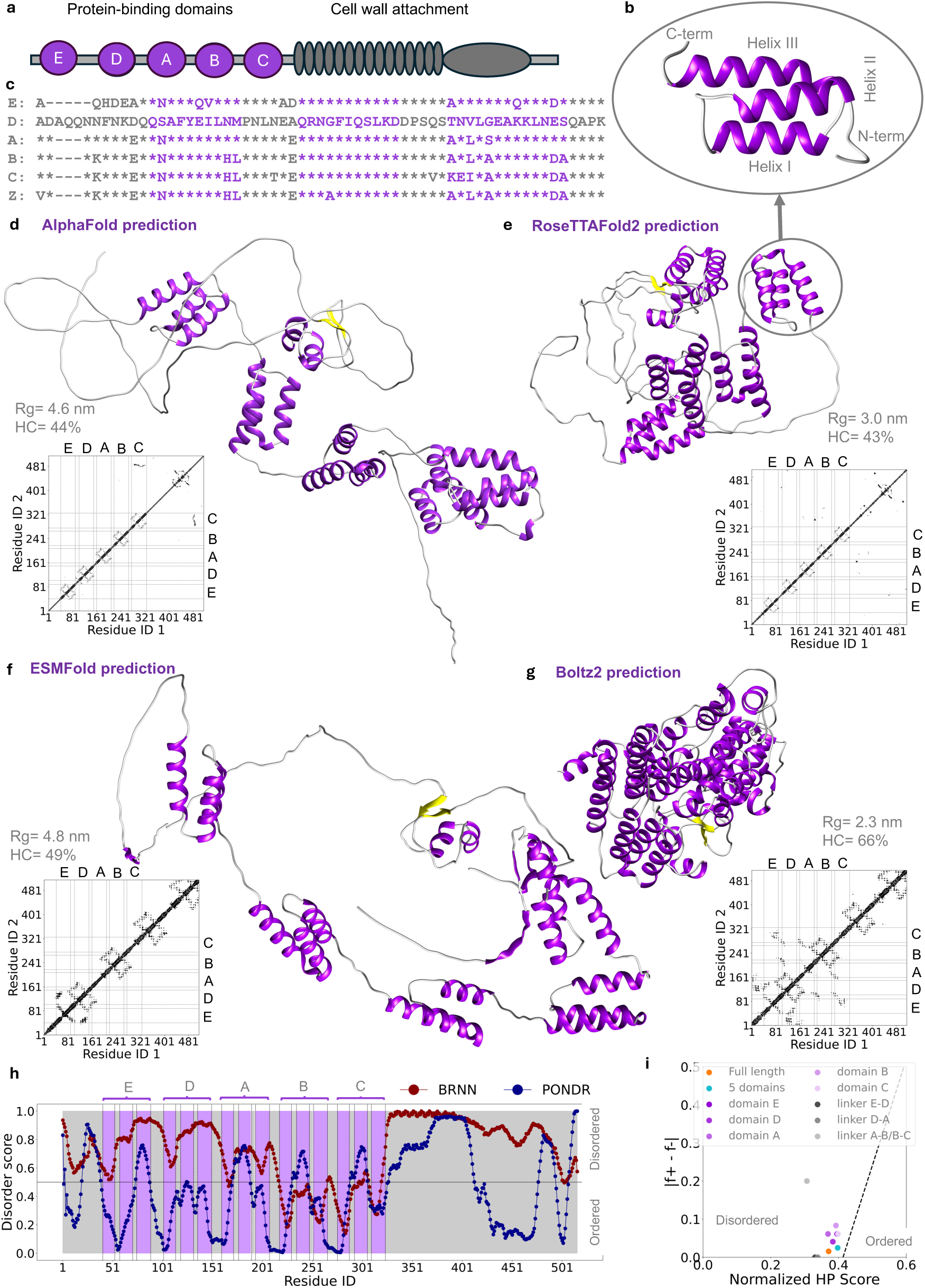
| Overview of the structural organization of full-length SpA. a, Schematic representation of the domain architecture of SpA. The five N-terminal IgG-binding domains (E, D, A, B, and C) are depicted as purple spheres and connected by flexible gray linkers. The C-terminal region, responsible for cell wall attachment, is shown in light gray. This illustration emphasizes the modular and flexible organization of SpA. b, The three-helix bundle structure of IgG-binding domains. c, Sequence alignment of individual domains and the engineered Z domain. AI-predicted structures for full-length SpA, with R_g_, helical content (HC), and all-atom contact map for d, AlphaFold^32^ e, RoseTTAFold2^33^ f, ESMFold^34^ and, g,, Boltz2.^35^ h, Predictions of structural order (scores < 0.5) and disorder (scores > 0.5) for full-length SpA calculated by the VL-XT algorithm (http://www.pondr.com/) and by Metapredict (predict disorder function, a bidirectional recurrent neural network^36^). i, Full-length SpA and its different regions mapped on the Uversky diagram.^37^

Widely used in biotechnology for antibody purification, each of the five tandem Ig-binding domains (and engineered analogs of these domains) of SpA is well-characterized. They fold independently into a stable three-helix bundle, and are also confidently predicted to be so by the AI algorithms. Yet the folding behavior of the full-length construct remains largely unexplored. Does the full-length protein fold as the sum of its parts? Experimental evidence hints otherwise. For instance, SpA’s full-length construct exhibits lower IgG-binding activity than its N-terminal half,^27^**^?^** and binding stoichiometry shifts from 2:1 (domain:IgG) to 1:1 in full-length.^28^ Small-angle X-ray scattering reveals unexpectedly compact dimensions,^2^ while surface behavior appears rigid rather than flexible.^29^ Collectively, these results point to behavior that cannot be explained by simply extrapolating from individual domains or relying on static predictions.

To address these questions, here we performed all-atom molecular dynamics simulations of full-length SpA in explicit solvent, using AI-predicted structures integrated into enhanced sampling to map the equilibrium folding free energy landscape. We complemented these simulations with experimental biophysical characterization, including circular dichroism spectroscopy and differential scanning calorimetry. Together, these analyses converge on a striking result: as opposed to predictions by AI models, full-length SpA does not fold into a stable three-dimensional structure in solution. Despite the individual domains’ known stability in isolation, none adopt their folded forms in the full-length context. Instead, the full-length protein remains compact yet predominantly structurally disordered, populating a heterogeneous ensemble with only residual secondary structure.

These findings challenge long-held assumptions about SpA’s modularity and highlight the importance of thermodynamic context in understanding multidomain protein behavior and the critical role of interdomain coupling. To the best of our knowledge, this is the first atomistic free energy landscape projection for a full-length multidomain protein of this size in explicit solvent, revealing that its domains fail to stabilize under native assembly. More broadly, our results show how thermodynamics-aware simulations can test and refine structure predictions—revealing hidden constraints on folding and unlocking new avenues for multidomain protein design.

## Results

### Predictions based on sequence

Fig. 1a illustrates the architecture of SpA schematically, highlighting its domain organization. SpA comprises five tandem Ig-binding domains (E, D, A, B, and C), each of which is known to adopt a compact three-helix bundle structure^4^ (Fig. 1b) separated by conserved short linkers. A sequence alignment of the domains and the engineered Z domain (a B-domain variant widely used in affinity purification) is shown in Fig. 1c. The structure predictions from four AI models for full-length SpA (516 residues) are compared in Fig. 1d–g along with the radius of gyration (*R_g_*), helical content (HC), and all-atom contact maps (6 Å cutoff) annotated with domain positions (E–D–A–B–C). Here, HC is calculated using the Dictionary of Secondary Structure on Proteins (DSSP)^30^ algorithm. The predicted structures colored by confidence are also presented in Extended Data Fig. 1c. Among these predictions, AlphaFold (Fig. 1d) and RoseTTAFold2 (Fig. 1e) predict structures expected from isolated domains in order, whereas ESMFold (Fig. 1f) and Boltz2 (Fig. 1g) merge segments around the E–D boundary into an apparent four-helix bundle and misplace other bundles relative to the sequence. The models also differ markedly in global compactness: AlphaFold (Fig. 1d)/ESMFold (Fig. 1f) yield more expanded conformations (*R_g_* ≈ 4.6–4.8 nm), while RoseTTAFold2(Fig. 1e)/Boltz2 (Fig. 1g) predict more compact states (R_g_ ≈ 3.0 and 2.3 nm, respectively). HC spans 43–66% (lowest in RoseTTAFold2 (Fig. 1e), highest in Boltz2 (Fig. 1g)). Residue-level DSSP (Extended Data Fig. 1a) shows that Boltz2 assigns extensive N-terminal helicity. In addition, heavy-atom contact maps with 2.5 Å cutoff showed unphysical contacts (steric clashes) for RoseTTAFold2, ESMFold, and Boltz2 (Extended Data Fig. 1b). Similar geometry issues have been reported by Haley *et al.*^31^

We also probed sequence-based disorder (Fig. 1h): PONDR score and BRNN predictor both show scores exceeding 0.5 encompassing Ig-binding domains. On the Uversky plot, the full-length protein, its N-terminal five-domain construct, linkers, and even the embedded domains in the full-length sequence cluster within the disordered regime (Fig. 1i). These divergent expanded vs compact folds, variable helicity, and disorder propensities motivated a thermodynamic analysis of SpA’s folding landscape.

### Destabilization of Ig-Binding Domain Folding in Full-Length Context

To assess the conformational behavior and thermal stability of full-length SpA, we performed exhaustive atomistic molecular dynamics simulations with enhanced sampling (parallel tempering in the well-tempered ensemble combined with well-tempered metadynamics [PTWTE-WTM] and multithermal-multiumbrella on-the-fly probability enhanced sampling [MM-OPES]) and experimental characterization.

After ensuring proper equilibration and convergence of the simulations (Extended Data Fig. 2-4, also see *Convergence* subsection of the *Methods* section), we computed the folding free energy landscapes (FEL) of SpA in solution and compared it with structural predictions from the AlphaFold and SpA domain structures available in the Protein Data Bank (PDB). We first did these FEL calculations at room temperature to resolve full-length SpA’s behavior in a room-temperature solution. We then extended these calculations to multiple temperatures to extract the thermal stability and compare it with the experimental data.

Although each of SpA’s five immunoglobulin-binding domains (E, D, A, B, and C) is known to fold stably in isolation,^38–42^ our findings showed that full-length SpA fails to stabilize structures with folded Ig-binding domains at room temperature equilibrium (Fig. 2a), as evident from the high free energy of the folded state (high Q-values) compared to the unfolded state (low Q-values), which is the minimum free energy state.

**Fig. 2.**
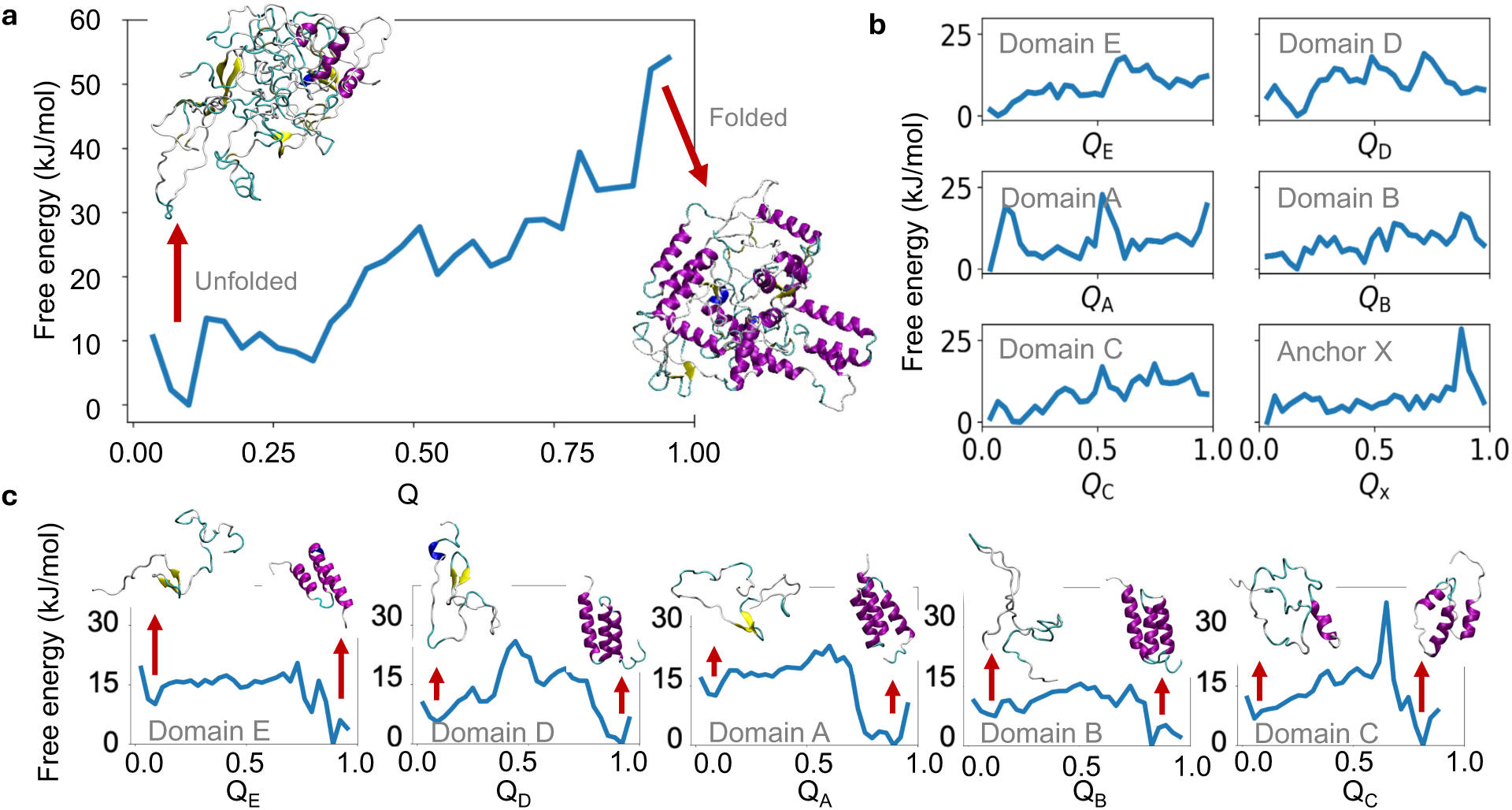
| Folding free energy landscape (FEL) of various SpA projected onto the fraction of native contacts (Q). a, FEL of full-length SpA shows no stable folded minimum. The free energy steadily increases toward high Q values, indicating that folded-like states are thermodynamically unfavorable. Representative structures from low-Q (unfolded-like) and high-Q (folded-like) regions are also shown. b, FEL of each domain within the full-length SpA context. None of the domains exhibits a stable minimum near Q = 1, indicating that their native folded states are not stabilized in the multidomain configuration. c, FEL of the same domains in isolation, each showing a clear minimum around Q=1, consistent with stable folding into their native structures.

We constructed the order parameter, *Q*, using the high and very high confidence parts (individual domains) in the AlphaFold predicted structure (Extended Data Fig. 1c) as the reference native structure. The one-dimensional free energy profile projected on the fraction of native contacts (*Q*) shows a monotonic increase toward high *Q* values, with conformations with folded Ig-binding domains (*Q* > 0.9) being over 40 kJ/mol higher in free energy compared to unfolded ones (*Q* < 0.1) (Fig. 2a). This shows that full-length SpA does not thermodynamically favor the formation of a stable, native-like conformation in the multidomain context.

To assess whether any of the five Ig-binding domains retain their folded conformation within the full-length SpA context, we calculated domain-specific *Q*. For this purpose, we redefined a *Q* for each of the five domains individually, and computed their respective free energy projections within the full-length construct (Fig. 2b). This analysis is particularly important because even though the full-length protein is not fully folded, some of the individual domains could still be folded within the full-length context. Strikingly, none of the domains showed a stable minimum near their *Q* = 1 (Fig. 2b), indicating that folded states are not stabilized even locally. This is in stark contrast to the folding behavior of the same domains in isolation, where each adopts a stable three-helix bundle (Fig. 2c). These findings suggest that interdomain interactions in the full-length sequence prevent even the formation of stable three-helix bundles in any domain (also see the *Contact maps* subsection).

Since the native contact definitions for *Q* were based on the AlphaFold-predicted structure of the full-length protein, we also assessed the reliability of the predicted structure for the individual domains with known crystal structures. We compared the structures of domains B and E (in the full-length) to their respective crystal structures deposited in the Protein Data Bank (PDB) (PDB: 1BDD, 1EDI) by calculating root-mean-square distance (RMSD) for backbone atoms between them. We found RMSD of 0.17 and 0.044 nm, respectively, which further confirms that the individual domain structures predicted by AlphaFold in the context of full-length protein fairly resemble the structures of isolated domains.

We also note that the simulations of isolated domains served as internal controls to validate the accuracy of the force field in this work. Using the same CHARMM36 force field and the same modified TIP3P water model, these simulations consistently predicted that each domain stably folds into its native three-helix bundle, exhibiting a free energy minimum near Q = 1 (Fig. 2c) at 300 K. These results reinforce the conclusion that the unfolded behavior of full-length SpA arises from its multidomain context.

To experimentally probe the thermal stability of SpA and validate our computational findings, we performed differential scanning calorimetry (DSC) on both the full-length protein and a single Ig-binding domain in isolation, domain Z. The DSC thermogram of full-length SpA (Fig. 3a) revealed a broad, low-amplitude, and mildly asymmetric transition, indicative of a heterogeneous and low-cooperativity melting process. In contrast, domain Z exhibited a sharp, well-defined peak, consistent with a highly cooperative unfolding transition. We will investigate the structural basis of the broad, asymmetric transition of the full-length SpA in the *Secondary structures* subsection.

**Fig. 3.**
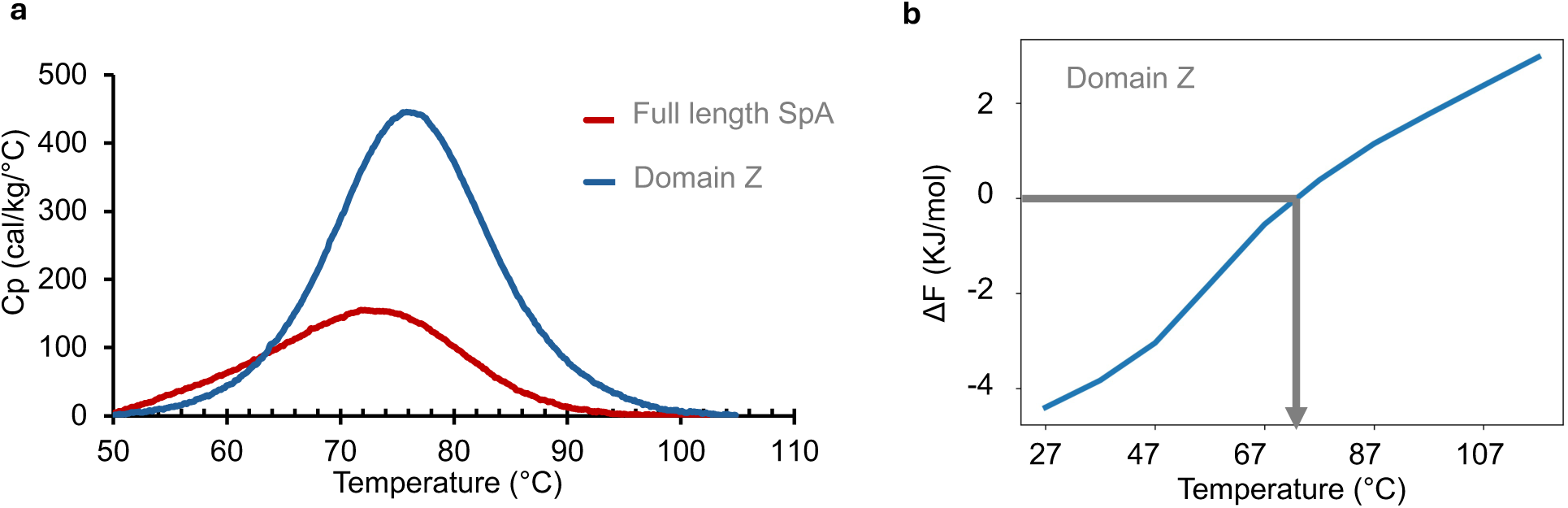
| Thermal stability of full-length SpA and isolated domain Z, both from computations and experiments. a, DSC thermograms for full-length SpA and domain Z. b, Estimated melting temperature from isolated domain Z simulation. The difference in the free energy of folded (F) and unfolded (U) states is calculated from FELs. The equation 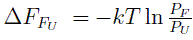 is used to calculate the stability difference between F and U states. *P_F_* and *P_U_* are unbiased probabilities of finding the system in these states, reweighted to the given range of temperatures. The melting temperature is estimated as the temperature where two probabilities are equal (shown with gray arrow), for domain Z.

To enable a direct comparison with experimental data, we quantified the thermal stability of domain Z using our FELs evaluated at multiple temperatures. Assuming a two-state model, we calculated the folding free energy difference (Δ*F*) between folded and unfolded states across a range of temperatures. The melting temperature (*T_m_*) was identified as the point where the two states are equally probable (see also ref.^43^) This analysis yielded a simulated *T_m_* of domain Z as 73*^○^*C (Fig. 3b), which is in excellent agreement with the DSC-derived value of 76*^○^*C (Fig. 3a, the peak of the blue data set).

Taken together, these results demonstrate that while isolated domains of SpA possess strong intrinsic foldability, the full-length protein does not support the formation of stable folded Ig-binding domains. The emergent picture of context-dependent instability prompts further structural analysis to pinpoint the molecular origins of this destabilization within the unfolded ensemble.

### Secondary structures

Here, we analyzed the secondary structures of SpA, focusing on turns and helices separately. To provide further reference for these analyses, we performed an additional control simulation termed “restricted-folded” full-length simulation. In this setup, the Ig-binding domains were restrained to their canonical three-helix bundle conformations, while the remainder of the protein (including N-terminus, linkers, and C-terminal anchor domain) remained flexible (See *Enhanced sampling methods* subsection of the *Methods* section).

We first examined turn formation using backbone dihedral angles to determine whether the required backbone conformations were properly formed for the turns. Previous studies have indicated that the turns connecting two α-helices play crucial roles in the folding kinetics of three-helix bundles and exhibit structural behavior similar to β-hairpin turns.^44^ Therefore, we analyzed the turn formation between helix 1 and helix 2 for each individual domain under three conditions for comparison: (i) main full-length simulation, (ii) restricted-folded full-length simulation, and (iii) individual domains simulated in isolation. The results for domain E in three conditions are shown in Fig. 4, and Extended Data Fig. 5 shows the same for each domain of the main full-length simulation.

**Fig. 4.**
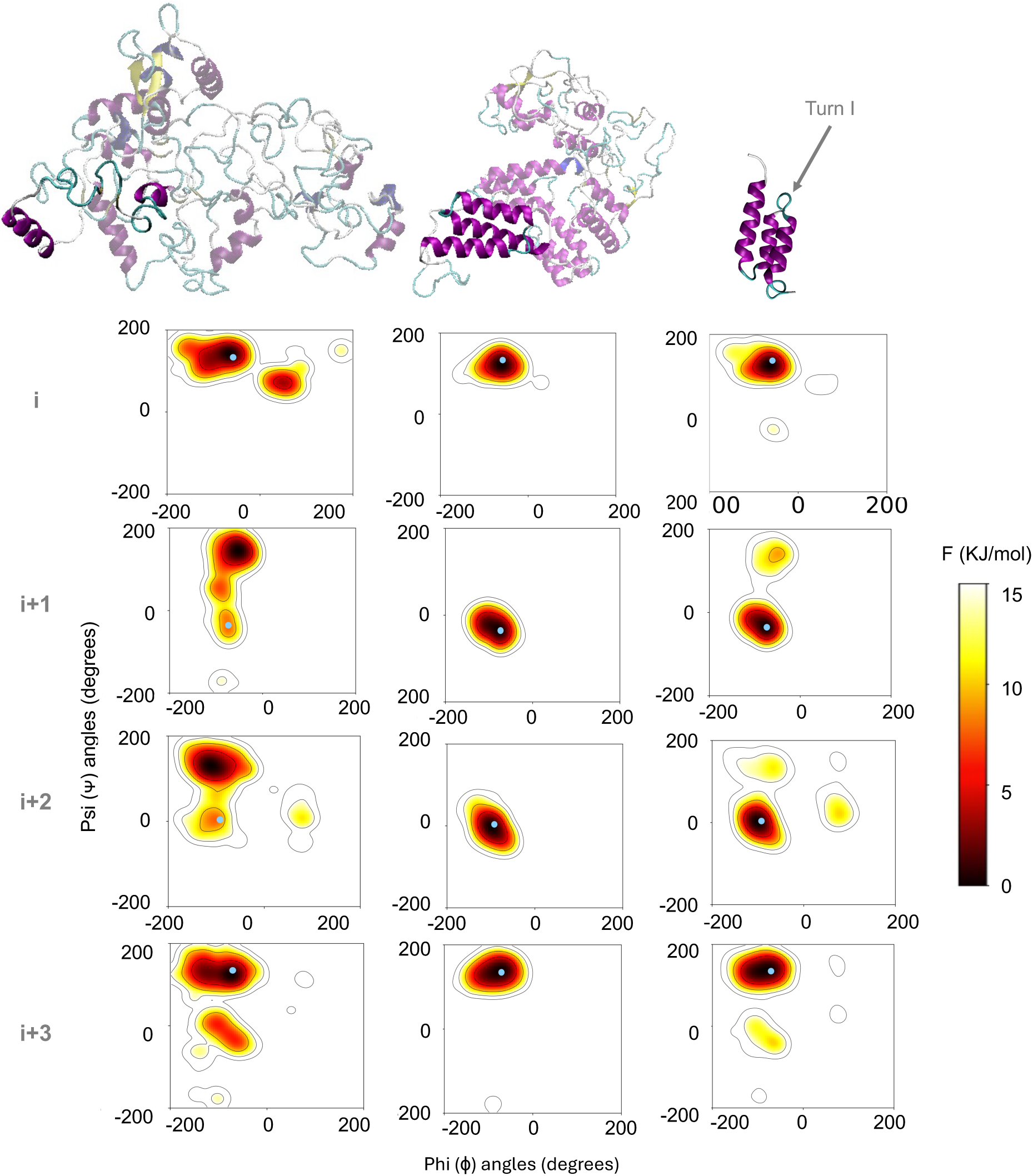
| Free energy surface projected onto dihedral angles for the residues of the first turn of domain E when it is in the. (1) main full-length simulation (left), (2) restricted-folded full-length simulation (middle), and (3) isolated domain simulation (right). These 2D free energy plots are calculated from the weighted histograms of dihedral angles. Dihedral angles of the amino acids at the equivalent position in the AlphaFold predicted structure are also shown as blue circles. The expected dihedral angles for the given turn are ≈ —60*^○^* – ≈ —30*^○^* for i +1 and ≈ —90*^○^* – ≈ —0*^○^* for i + 2, which is a type I turn. A representative snapshot of the most populated conformations from each simulation is presented on the left side of each panel.

The most populated angles for the turn residues *i* + 1 and *i* + 2 are within the vicinity of the expected *β*-turn type (type I) in both the isolated domain E simulation and restricted-folded full-length simulation (Fig. 4 right and middle, respectively), while these conformations were rarely sampled in the main full-length simulation (Fig. 4, left), which clearly shows that proper turn formation in the main full-length simulation is absent in the individual domains (where the triple-α-helical structures are expected) when they are in the full-length assembly.

We next analyzed the helix formation in this protein, which constitutes the defining feature of the immunoglobulin-binding domains of SpA. Prior studies have shown that the folding of these three-helix bundles occurs cooperatively, typically nucleating around helix II, followed by packing of helices I and III^45,46^). Recent simulations support this mechanism, revealing correlated formation of helices II and III, and lower stability of helix I.^47^ In addition, Myers and Oas demonstrated that preorganized helices in the unfolded ensemble—particularly helix III—facilitate rapid folding through diffusion–collision dynamics.^48^

To characterize the extent of helical content in full-length SpA, we calculated helix propensities across simulation (sub)trajectories using the DSSP algorithm (Fig. 5a). As expected, high helix propensities were observed in the folded subensemble (Q > 0.9) of the main full-length simulation and in the restricted-folded simulation, where the Ig-binding domains were artificially restrained into triple-helix bundles (for the restricted-folded simulation). However, in the equilibrium ensemble of the full-length construct, the same regions exhibited markedly lower helicity. This suggests that, although some local helices can form partially and transiently in the full-length context, the cooperative packing of three-helix bundles is largely disrupted.

**Fig. 5.**
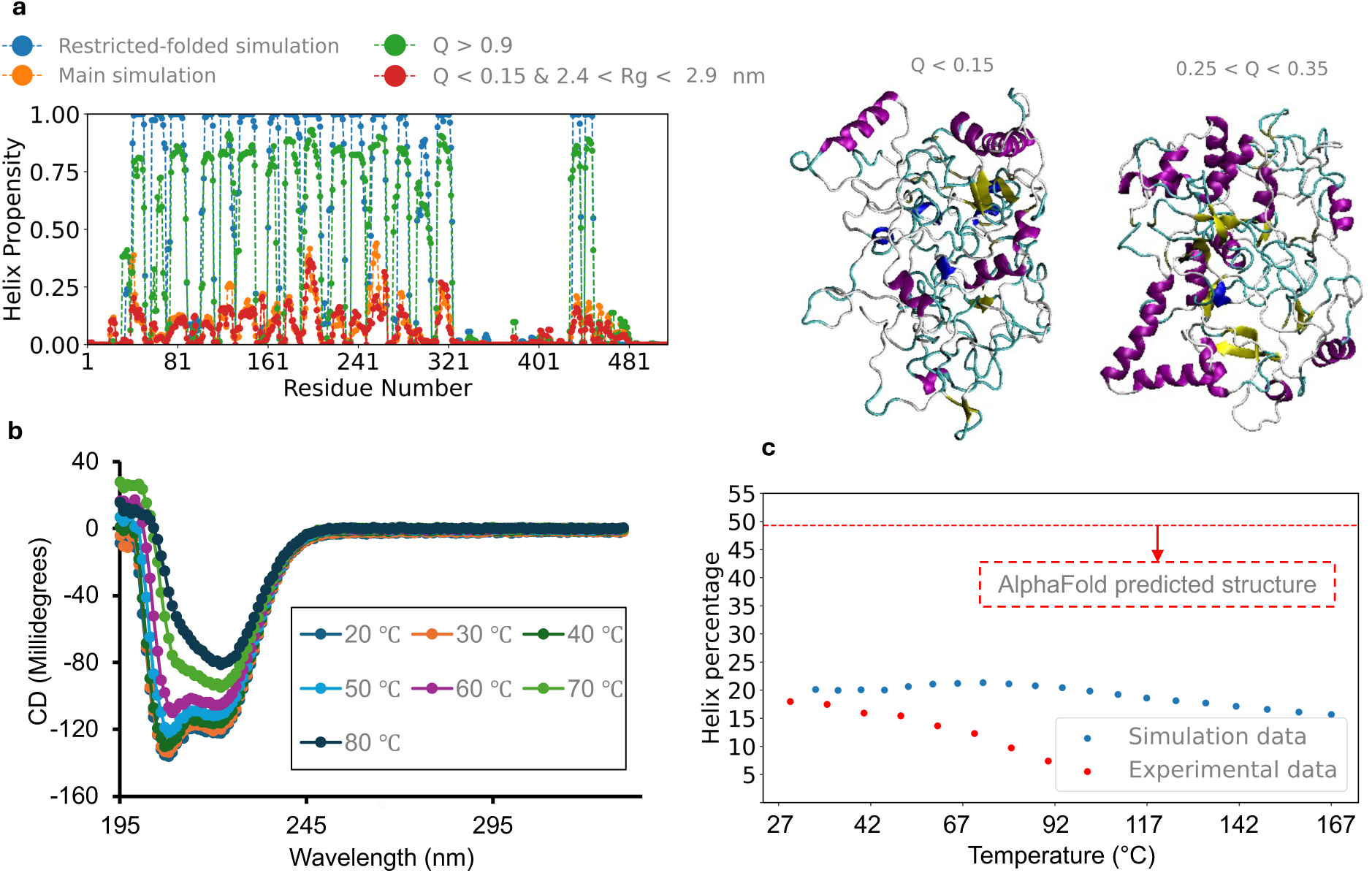
| Computational and experimental characterization of helices of Protein A. a, Helix propensities of full-length SpA calculated from DSSP analyses of restricted-folded simulation, main simulation, and two (sub)trajectories of the main simulation. b, CD spectra of full-length SpA c, Helix percentage of full-length SpA calculated from CD spectroscopy and the simulation.

This incomplete helix formation is also consistent with the broad, asymmetric, and low-intensity DSC peak observed for full-length SpA (Fig. 3a) in the sense that the broad, asymmetric peak corresponds to gradual melting of partial helices. To connect structural features with thermodynamic signatures, we calculated the change in helical content (HC)–weighted heat capacity (Cp × HC) from the simulation data. The resulting curve (Extended Data Fig. 6b) revealed a diffuse thermal transition, reinforcing the interpretation that thermal unfolding in full-length SpA reflects gradual loss of partial helices, rather than a sharp cooperative unfolding event. In contrast, domain Z displayed a well-defined, narrow transition in both simulation and experiment, characteristic of two-state folding.

As an experimental validation, (Fig. 5b) shows the results of circular dichroism (CD) spectroscopy to experimentally quantify HC across temperature. We presented the HC calculated from CD together with the HC from simulations (extracted from DSSP analysis across 20 temperatures). Both of them showed a gradual decrease with increasing temperature, mirroring each other and verifying the lack of a cooperative melting (Fig. 5c). Notably, the room-temperature HC in simulations was already significantly lower than that of the AlphaFold-predicted structure (Fig. 1d).

Visualizing the secondary structure over time further supports this picture (Extended Data Figs. 7–11). In the restricted-folded simulation, fifteen continuous helical segments are clearly visible, corresponding to the five structured domains (Extended Data Fig. 8). In contrast, the unrestrained full-length simulation shows only scattered, short-lived helices (Extended Data Fig. 7), localized to the same regions as in the folded form. This pattern suggests a degree of intrinsic helical preorganization within the unfolded ensemble, consistent with the notion proposed by Myers and Oas.^48^

Importantly, neither simulation-derived data nor experimental CD spectra (Fig. 5b) exhibited a sigmoidal temperature dependence for full-length SpA. This lack of a sharp transition further supports the absence of a cooperative, globular unfolding behavior. In contrast, the same DSSP technique applied to domain Z showed a clear sigmoidal transition in HC over temperature (Extended Data Fig. 6a), highlighting the key difference in folding/unfolding behavior between isolated domains and their assembly in the full-length constructs. The broad and gradual decrease in ellipticity likely reflects multiple, partially overlapping transitions among domains and linkers, which are not individually resolved in the 222 nm signal.

Together, these results reveal that the full-length SpA ensemble is dominated by partially structured states with residual helical character. This ensemble-level heterogeneity underlies the distributed unfolding behavior observed thermodynamically and highlights the failure of modular folding in this context. In the following section, we investigate potential sequence-encoded or interdomain features that may underlie this disrupted folding behavior.

### Conformational Properties Reveal Globular Yet Unfolded SpA

Since the fully-folded conformation of full-length SpA is not stable, we sought to characterize its global structure and compactness using polymeric descriptors such as radius of gyration (*R_g_*), and internal polymer scaling exponents.

The free energy projection here reflects the weighted joint distribution of *R_g_* and Q (Fig. 6). The lower free energies correspond to higher probabilities. By projecting the free energy onto these two variables, we can simultaneously assess the compactness and folding level of full-length SpA.

**Fig. 6.**
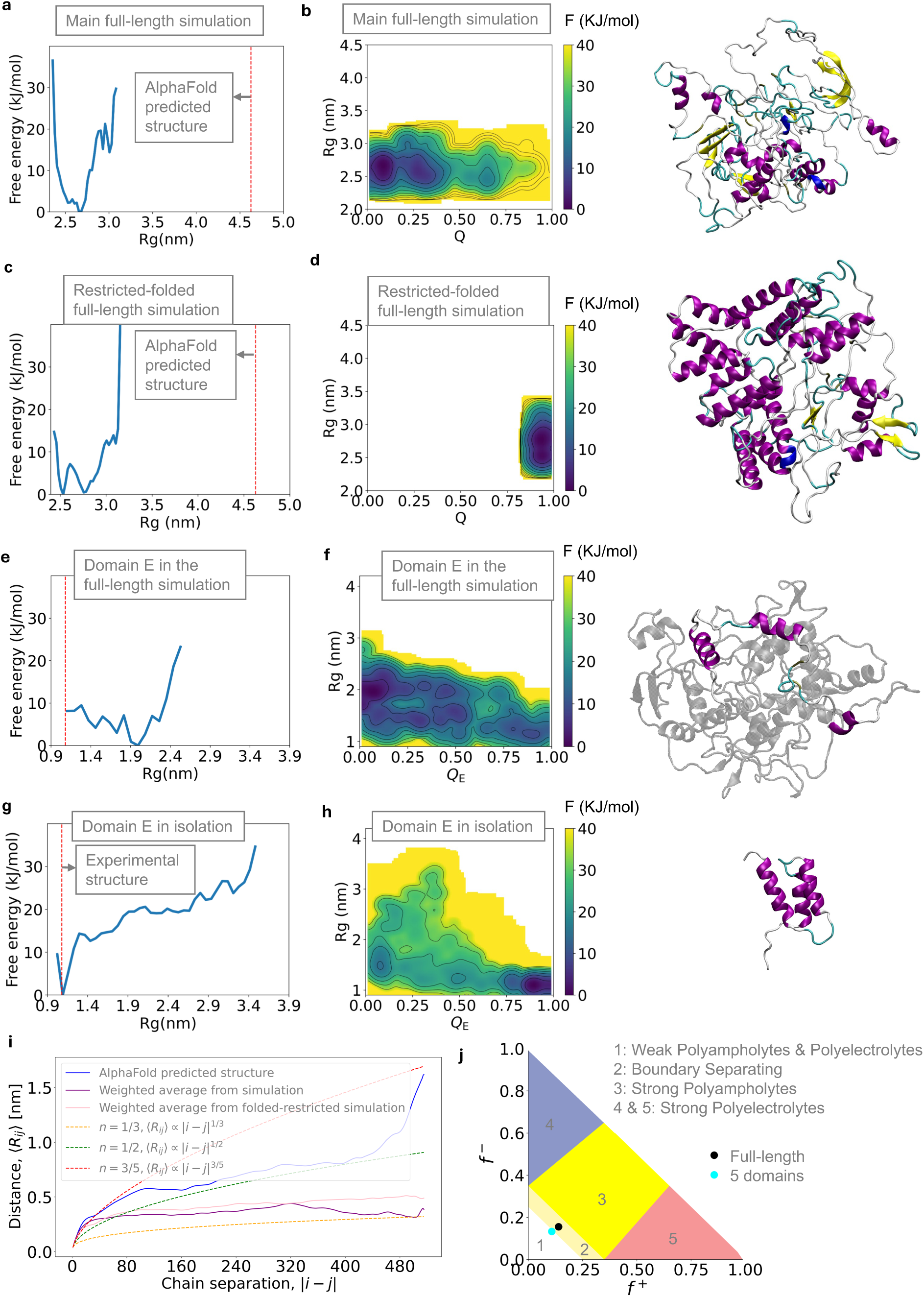
| Global and domain-level size projections. a, Free energy for the main full-length simulation projected onto *R_g_*. b, Free energy for the main full-length simulation projected onto Q and *R_g_*. c, Free energy for the restricted-folded full-length simulation projected onto *R_g_*. d, Free energy for the restricted-folded full-length simulation projected onto Q and R_g_. e, Free energy for the main full-length simulation projected onto *R_g_* of domain E. f, Free energy for the main full-length simulation projected onto R_g_ and Q of domain E. g, Free energy for the isolated domain E simulation projected onto *R_g_*. h, Free energy for the isolated domain E simulation projected onto Q and *R_g_*. i, Weighted ensemble-averaged interchain distance (< *R_ij_* >) for full-length SpA with respect to chain separation. j, Das-Pappu diagram for full-length SpA and 5 domains (excluding the C-terminal anchor domain).

Across the full-length simulation, *R_g_* remained narrowly distributed below 3 nm, reflecting a very compact ensemble of configurations (Fig. 6a) compared to the AlphaFold predicted structure’s *R_g_* (larger than 4.5 nm). A two-dimensional free energy projection onto *R_g_* and folding progress variable Q (Fig. 6b) revealed that compactness persists across a broad range of folding states, indicating that neither domain-level folding nor global structure formation is necessary to obtain this compact ensemble. Surprisingly, even the restrained-folded simulation yielded an ensemble that is nearly equally compact as the main full-length simulation (Fig. 6c) with five completely folded domains (Fig. 6d), suggesting that the full-length sequence favors compactness even when the individual domains are forced to retain their triple-helical structures.

To assess whether the sizes of individual domains are affected by their incorporation into the full-length assembly, we projected the free energy landscape obtained from the main full-length SpA onto structural parameters specific to domain E, as shown in Figs. 6e and 6f. Here, *R_g_* is calculated using only the backbone atoms of domain E, and Q is the redefined fraction of native-like contacts based on atom pairs within this domain (*Q_E_*). For comparison, we also analyzed simulations of the isolated domain E, presented in Figs. 6g and 6h. In the isolated simulation, the unweighted free energy projected onto *R_g_* reveals an absolute minimum at near 1 nm, and the two-dimensional surface projected onto *R_g_* and *Q_E_* (Fig. 6h) confirms that this basin corresponds to a folded state with high values of *Q_E_*. The *R_g_* of the experimentally determined solution NMR structure of domain E (PDB ID: 1EDI) is also indicated by the dashed red line, which aligns with the *R_g_* of the folded state observed in simulation.

Notably, domain E within the full-length SpA exhibits a slightly higher *R_g_* compared to its fully-folded state in isolation. This subtle expansion suggests that, although the overall assembly of full-length SpA is highly compact, individual domains remain somewhat loosened, possibly due to incomplete turn formation (Fig. 4). Such partial unfolding could facilitate transient inter-domain interactions, a feature further examined in the following section.

We also evaluated the balance of protein-solvent, protein-protein, and solvent-solvent interactions, which determine the overall dimensions of polymeric materials, like proteins. Also known as polymer scaling laws, internal distance scalings provide a quantitative framework for describing how internal distances scale with chain length, distinguishing between swollen coils, ideal chains, and compact globules.^49–51^ Therefore, to further investigate the structural organization of full-length SpA, we examined the ensemble-averaged internal distances and applied scaling laws to characterize its behavior.

We used the scaling relation < *R_ij_* > = *A|i* — *j|^v^* for the weighted ensemble-averaged internal distance, < *R_ij_* >, as a function of chain separation, *|i* — *j|*. Following our previous work,^50^ we applied a global prefactor of *A* = 0.55 nm and plotted limits with three different exponents. The limits of ⟨*R_ij_*⟩ ∝ *|i — j|*^0.6^, ⟨*R_ij_*⟩ ∝ *|i — j|*^0.5^, and ⟨*R_ij_*⟩ ∝ |i — j|^0.33^ describe how monomer interactions influence chain expansion or compaction. A scaling exponent of *v* = 0.6 corresponds to a swollen coil with effectively repulsive interactions, *v* = 0.5 represents an ideal chain with balanced interactions, and *v* = 0.33 describes a globular state with effectively attractive interactions.^49,50^

Fitting the above equation to our data, we obtained exponents of 0.11, 0.22, and 0.48 for the main simulation, restricted-folded simulation, and AlphaFold predicted structure, respectively (Fig. 6i). When we limited the analysis to shorter chain separations (up to 80 residues), the exponents increased to 0.29, 0.4, and 0.45, respectively (Extended Data Fig. 12). This shift suggests the presence of blob-like structural organization^49,52^ within SpA, where local segments follow a more expanded conformation before transitioning into a compact state at larger distances. This local looseness aligns with the slightly elevated R_g_of domain E (Fig. 6e), reinforcing that the compactness of full-length SpA coexists with modest domain-level reduced packing.

Overall, the low global exponent confirms that the full-length SpA adopts very compact globular conformations that do not scale after a certain chain separation, i.e., reach a plateau (Fig. 6i). Of note, the full-length SpA falls into region 2 in the Das-Pappu diagram,^53^ the boundary separating 1 & 3, supporting its classification as a globular protein (Fig. 6j). It should be noted that the term “globular” in this case does not indicate a folded protein, but an unstructured, globe-like compact protein.

It is worth mentioning that we calculated the average of root mean square fluctuations (RMSF) for backbone atoms in the isolated domain E simulations (where the protein actually folds) as 2 nm, whereas RMSF decreased to 1.4 nm in the main full-length simulations. This reduction implies that the compactness of the full-length SpA restricts local fluctuations, preventing each domain’s triple-α-helical structure from folding independently. To further investigate the underlying causes of this compactness, we will analyze contacts between different amino acids in the next session.

### Atomistic Contact-Level Evidence Related to Interdomain Compaction

The compactness of full-length SpA (Fig. 6) arises despite the absence of domain-level folding, raising the question of what types of residue–residue interactions underlie this collapse. To address this, we computed residue–residue contact maps (averaged over all heavy atoms) for four ensembles: i) the AlphaFold-predicted structure (Fig. 7a), ii) the restricted-folded simulation (where domains are held folded) (Fig. 7b), iii) the folded subtrajectory (*Q* > 0.9) from the main simulation (Fig. 7c), and the unfolded subtrajectory (*Q* < 0.15) from the main simulation (Fig. 7d). Each contact map shows the fraction of times that any given pair of residues is found within 6 Å of each other, except the AlphaFold contact map. Since it represents a single structure, instead of an ensemble-average, it is shown as a binary contact map.

**Fig. 7.**
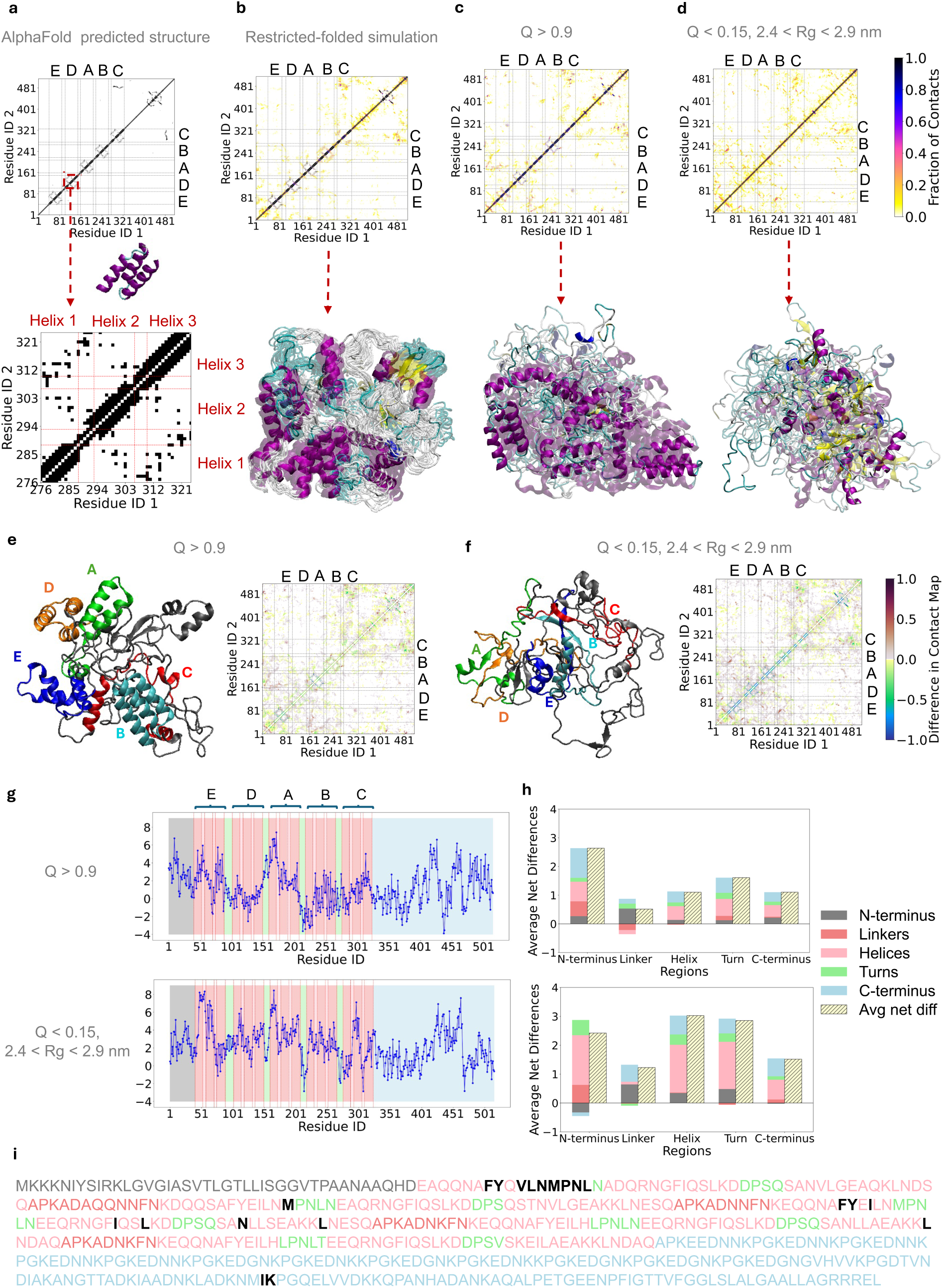
| Contact map for. a, AlphaFold predicted structure b, restricted-folded simulation c, the main simulation folded subtrajectory (*Q* > 0.9) d, the unfolded subtrajectory (*Q* < 0.15, 2.4 < *R_g_* < 2.9*nm*). Difference in contact fractions obtained by subtracting the subtrajectory contact probabilities from the restricted-folded simulation for e, folded f, unfolded. g, Sum of the contact map differences after excluding the contacts presented in the AlphaFold structure, plotted per residue. h, Average of these summed differences across distinct regions of SpA. i, Full-length SpA sequence highlighting residues with total contact difference > 6 in the unfolded subtrajectory (black).

In the contact map of the AlphaFold-predicted structure, five distinct rectangles near the diagonal correspond to the five triple-α-helical domains, as highlighted for domain D (Fig. 7a). These blocks exhibit characteristic patterns: diagonal lines for intra-helix contacts within a single helical segment, and dense off-diagonal features for helix-helix packing (for three-helix bundle architecture).

In the restricted-folded ensemble (Fig. 7b) and *Q* > 0.9 ensemble (Fig. 7c) the off-diagonal features characteristic of helix–helix packing are slightly diminished for some domains, reflecting a partial loss of bundling. In addition, both contact maps display numerous *unpredicted contacts* absent in the AlphaFold structure (Figs. 7b-c), which contribute to the compactness of this protein(Figs. 6a-d). The number of these *unpredicted contacts* increases as *Q* decreases, indicating the accumulation of unpredicted interactions in more unfolded configurations. Collectively, these results suggest that our physics-based simulations produce both short-range and long-range residue-residue contacts for the full-length SpA that are not reproduced by the AI-based predictions.

Since the unfolded state of full-length SpA encompasses the most probable configurations (Fig. 6b), it is important to identify the contacts that are specifically formed in this unfolded ensemble but absent in the fully-folded structure. To quantify these differences, we computed contact map differences by subtracting the contact map of each subtrajectory from that of the restricted-folded reference (Figs. 7e and f). Blue regions indicate contacts lost relative to the folded reference (typically intra-domain), while brown regions indicate new contacts gained (typically interdomain). The unfolded subensemble (*Q* < 0.15) shows a clear loss of intra-domain contacts (corresponding to the triple-α-helical domains) and a marked gain of interdomain contacts, particularly in regions spanning domains E through C, that is, residues < 327 (Fig. 7f).

We next sought to determine which residues are most involved in these newly formed interdomain contacts. To do this, we excluded contacts already present in the AlphaFold model and calculated the net gain or loss in contact probability for each residue across the trajectory (Fig. 7g). Residues within the triple-helix regions (pink regions) show the highest accumulation of unfolding-induced contacts in the unfolded ensemble.

To distinguish the contributions from different regions, we grouped residues into five categories: N-terminus, Linkers, Helices, Turns, and C-terminus. We then calculated the average contact change per residue in each region (Fig. 7h). Here, the *N-terminus* refers to residues preceding domain E; the *C-terminus* to residues following domain C; *Linkers* are the linker regions between Ig-binding domains; and *Helices* and *Turns*, are the helix and turn regions in the Ig-binding domains, respectively. The Helices region (pink) showed the largest increase in unpredicted contacts, followed by the Turns (green), suggesting that helical cores from different domains interact extensively upon unfolding. These results directly explain the loss of independent domain folding in full-length SpA.

Finally, we mapped residues with the highest contact differences (cumulative sum of *unpredicted contacts* (Fig. 7g) greater than 6) in the unfolded ensemble onto the full-length sequence (Fig. 7i). The sequence of full-length SpA is color-coded according to these structural regions (Fig. 7i). Amino acids with a cumulative sum of *unpredicted contacts* (Fig. 7g) greater than 6 in the unfolded ensemble are highlighted in black. Sequence analysis revealed that more than 75% of residues with the highest *unpredicted contacts* (> 6) are hydrophobic, consistent with the observation that helical regions, which are predominantly hydrophobic, contribute most significantly to the emergence of these new contacts in the unfolded state (Fig. 7h).

## Discussion

Our results revealed that full-length SpA does not adopt stable domain folds in solution, despite each Ig-binding domain folding in isolation and being confidently predicted by sequence-based AI models. This challenges the common modular view of multidomain proteins—an area increasingly recognized as a blind spot in folding studies. ^8,18,54^ Rather than treating structure as a single static model, we characterized the full-length SpA as an ensemble using enhanced-sampling atomistic simulations and corroborating biophysical measurements (CD and DSC), providing an explicit solution-state view.

Our simulations predicted that, in the absence of ligands, full-length SpA adopts a compact conformation with only partial helices, rather than the five neat triple-helix bundles predicted by AI models (Fig. 1d-g). This finding was surprising, as the same simulation protocol correctly predicted each isolated three-helical bundle as a two-state folding protein. Comparisons of melting temperatures and R_g_ values for the isolated domains with previous experimental measurements showed good agreement, reinforcing the reliability of the simulation approach. Additionally, experimental techniques including DSC and CD spectroscopy were employed to characterize full-length SpA, ruling out the possibility that the compact unfolded ensemble was simply a simulation artifact.

Analysis of the backbone dihedral angle distributions indicated a lack of readily formed turns in full-length SpA (Figs. 4 and Extended Data Fig. 5), while the isolated Ig-binding domains readily formed turns, suggesting that turn formation, a critical step for triple-helix bundle folding, is inhibited in full-length assembly. Furthermore, CD spectroscopy and DSSP analysis from simulations showed that the extent of helicity in full-length SpA was less than half that predicted by AlphaFold (Fig. 5c). The scanning calorimetry results indicated non-cooperative melting, as evidenced by a broad and asymmetric DSC peak (Fig. 3a). Via CD (experiments) and DSSP (simulations) calculations, we showed that this non-cooperative melting is consistent with the melting of partially formed helices. Although partial helices are present even in the unfolded ensemble of full-length SpA (Fig. 5a), the long-range contacts within each domain that would stabilize them into a cooperative fold are missing, leaving domains unable to achieve a stable three-helix bundle. Myers and Oas observed that the isolated B domain also exhibits nascent helices in its unfolded state, which they termed “preorganized secondary structure”.^48^ Our results indicate that while partial helices are also present in full-length SpA, turn initiation and stabilizing long-range contacts for each domain are both suppressed, preventing the formation of a stable fold. Therefore, instead of forming a globular triple-helix bundle, each bundle extends to have contacts with other domains (Fig. 6e and Fig. 7g).

To understand why the otherwise stable isolated domains become destabilized in the full-length assembly, we further analyzed the protein’s conformational properties and inter-residue contact patterns. R_g_, and polymer scaling analyses revealed that full-length SpA is highly compact. Contact map analyses showed that this compactness arises primarily from inter-domain interactions, particularly among hydrophobic residues. Given the high sequence identity (75%) between SpA domains,^55^ such inter-domain interactions (such as domain swapping attempts) are plausible. This observation is consistent with previous findings stating that the interaction between neighbor domains with similar sequence identity is highly probable.^15,56,57^

Among the AI models we evaluated, none accurately captured the secondary structure of full-length SpA. However, RoseTTAFold2 produced the most consistent single-structure prediction relative to our physics-based simulations and experiments, exhibiting a more compact arrangement (lower R_g_) and reduced HC. Indeed, RoseTTAFold2 was able to recover a subset of long-range contacts between domains absent in the AlphaFold output, named “unpredicted contacts” here. Although both models benefit from multiple sequence alignments (MSAs), the improved agreement likely reflects architectural priors. In particular, RoseTTAFold’s three-track design jointly reasons over sequence (1D), pairwise/distance maps (2D), and coordinates (3D), which may encourage weak but physically more plausible inter-domain packing.^33^

More broadly, AI predictors typically output one structure with a confidence score; they are not designed to return thermodynamically weighted ensembles or conformational exchanges, which limits their realism for systems that populate multiple states. This “single-snapshot” limitation is widely noted and contributes to discrepancies for flexible, multidomain, or disordered proteins.^58–61^ We also argue that the divergence between AI models and our simulations is related to a shared training bias: all predictors ultimately derive structural supervision from the Protein Data Bank (PDB) or distillations thereof. As a consequence, these models are strongest on well-represented, well-folded domains. However, they struggle with multidomain proteins like full-length SpA that are underrepresented in PDB structure repository. This limitation, referred to as a “blind spot” of structure prediction algorithms, has been highlighted by Chakravarty *et al.*,^62^ and also reported by others.^63,64^ Therefore, while AI models for protein structure prediction are undoubtedly attractive in terms of reduced computational cost and time, their results should be interpreted with caution.

Molecular dynamics (MD) simulations, though computationally more demanding, generate ensembles governed by physics-based force fields and thermodynamics, enabling them to capture behaviors more consistent with experimental observations, even for systems outside the traditional structured protein space. MD simulations also allow us to assess how intrinsic crowding within multi-domain assemblies influences their thermodynamic landscape—an effect that is typically invisible to static AI predictions but likely important for understanding protein behavior.

Given that full-length SpA includes five homologous domains and a disordered C-terminus, it is plausible that even a reduced number of interacting domains could induce similar folding instability. Thus, identifying the minimum number of domains required to destabilize folding is an important future direction. Moreover, while SpA serves as an important case study, it cannot be assumed to be representative of all multidomain proteins. A broader, systematic investigation into how domain-domain interactions modulate folding pathways across diverse multidomain systems is also warranted.

Notably, SpA must traverse the membrane via the Sec secretion pathway,^65^ and we argue that the intrinsic unfolded tendency revealed here may be biologically advantageous, facilitating secretion and later refolding upon cell-surface anchoring. Another important future direction involves characterizing the conformational behavior of full-length SpA in the presence of its natural binding partners as well as when it is anchored on a surface. Protein secondary structure is often remodeled upon ligand binding, ^66–68^ and recent large-scale generative modeling studies have reinforced that conformational changes—including domain motions, local unfolding transitions, and the formation of cryptic binding pockets—are central to protein function and frequently coupled to binding events.^69^ In this context, it is plausible that the folded state could stabilize upon interaction with immunoglobulins or other physiological partners. Notably, these partial helices coincide with the Helices I, II, and III known to mediate partners’ binding.^26^ Therefore, we hypothesize that the partial helices observed even within unfolded conformations (Fig. 5b) may serve as initial recognition motifs, priming SpA for functional interactions—a possibility that merits further experimental and computational investigation.

## Methods

### AI-based Structure Predictions

The sequence of full-length SpA was obtained from UniProt P02976. We used https://alphafold.ebi.ac.uk and https://forge.evolutionaryscale.ai for structure predictions of AlphaFold and ESMFold for this given sequence, respectively. Predictions from RoseTTAFold2 and Boltz2 were obtained using their respective GitHub repositories at https://github.com/uw-ipd/RoseTTAFold2 and https://github.com/jwohlwend/ boltz, respectively. To assess the intrinsic disorder, we calculated disorder scores using the VLXT algorithm available at http://www.pondr.com/PONDR and the bidirectional recurrent neural network (BRNN) predictor implemented in the Python package Metapredict (predict disorder function).

### Modeling

We simulated the full-length SpA under two conditions: i) the main simulations, where all domains can fold or unfold freely (no restraints), and ii) restricted-folded simulations, in which the individual domains were restrained in their folded states (further details in the *Enhanced Sampling Methods* subsection). Additionally, we conducted separate simulations for each individual SpA domain in isolation to calculate their folding free energy landscapes. Since no experimentally determined structure was available, we used the AlphaFold-predicted structure (AF-P02976-F1-v2) as the initial configuration for the full-length SpA simulations. For isolated domain simulations, we used experimentally resolved structures when available and AlphaFold-predicted structures otherwise; PBD ID 1EDI for domain E, AlphaFold-predicted structures for domains A and D, PDB ID 1BDD for domain B, and PDB ID 4NPE for domain C.

The initial protein structures were placed in a truncated octahedron box with a volume of 1936 nm^3^ for full-length SpA and 180-280 nm^3^ for isolated domains. All systems were modeled using the CHARMM36 force field, combined with the modified TIP3P water model.^70^ Na^+^ ions were added to ensure electroneutrality in the full-length simulations. For isolated domain simulations, the salt concentration was adjusted to 50 mM (NaCl) to allow a fair comparison with experimental melting temperature measurements.

### Enhanced sampling methods

Parallel-tempering in the well-tempered ensemble combined with well-tempered metadynamics (PTWTE-WTM)^71–74^ was used for sampling the full-length SpA in the main simulations. In the framework of the well-tempered ensemble,^71^ the potential energy was used as the collective variable in well-tempered metadynamics using Gaussian kernels with a 1000 kJ/mol width and a 4 kJ/mol initial height. The deposition stride for the Gaussians was set to 1000 steps, and a bias factor of 50 was employed. The temperatures of the 20 replicas, ranging between 300 K and 440 K, were distributed geometrically.

In the WTM part of PTWTE-WTM, we biased a similarity-based order parameter (*Q*), which is defined as a fraction of native-like contacts, that is, a number defined between 0 and 1. The order parameter Q evaluates whether the domains of SpA adopt their folded conformations relative to a reference structure (that we deemed “native” structure). Since no crystallographic structure of full-length SpA is available in the literature, we used the AlphaFold-predicted structure as the reference for computing *Q*. It is important to note that the definition of Q is based solely on 5 folded domains, E, D, A, B, C, and a small segment of the cell wall anchor (domain X), (residues 422–466) that AlphaFold predicted to contain secondary structures. Consequently, linkers, turns, and other disordered regions were excluded from the definition of *Q*. The order parameter *Q* is defined as 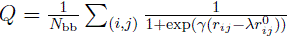 following the generalized definition given in references.^75,76^ The sum runs over N_bb_pairs, which is the total number of atomic pairs (*i, j*) that are considered in contact in the reference contact list. A heavy (non-hydrogen) backbone (bb) atom of a residue was considered to be in contact with a bb atom of another residue if the distance between them is less than 5 Å in the reference structure. The heavy bb atoms are considered as α-carbon, carbonyl carbon, carbonyl oxygen, and amide nitrogen. Here, r^0^ and r_ij_ are the distances between i and j in the reference structure and in any given instantaneous configuration, respectively. γ in the smoothing function was taken as 50 nm*^—^*^1^ and the adjustable parameter λ was taken as 1.5.^76^ For further details of the order parameter, we refer to our previous work.^77,78^

The initial Gaussian height for Q was set to 2 kJ/mol with a bias factor of 40 for the WTM sampling in PTWTE-WTM simulations. The Gaussian width was set to 0.01. Since Q is strictly defined between 0 and 1, interval limits were applied alongside restraining potentials to avoid accumulating systematic errors at the boundaries of Q (at Q = 0.01 and 0.99 for lower and upper boundaries, respectively).^79^

For the folded-restricted full-length SpA, we employed PTWTE with the same simulation parameters in the main full-length simulations. To restrain domains in their folded states, we applied a lower boundary constraint at Q = 0.95, using the same definition of Q described above.

The multithermal-multiumbrella on-the-fly probability enhanced sampling (MM-OPES)^80–83^ technique was applied for isolated domain simulations. The target distribution that MM-OPES aims to sample for our case is:

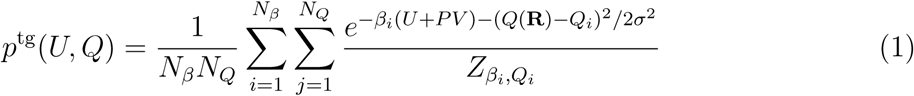

where the potential energy *U* and the number of native contacts Q are the collective variables, *β_i_* = 1/(*k_B_T_i_*) are inverse temperatures, *Q_i_*are the centers of the umbrella potentials, *P* is the pressure, *V* is the volume, and *Z_β__i,Qi_* is an appropriate partition function. We have used the notation *Q*(R) to highlight the dependence of *Q* on the atomic coordinates R. Once the minimum temperature *T_min_* = 1/(*k_B_β_N__β_*) and the maximum temperature *T_max_* = 1/(*k_B_β_1_*) are specified, intermediate temperatures, T_i_, are spaced geometrically. The number of *β_i_, N_β_*, is then automatically determined (see^80^ for further details). The umbrella potentials were equispaced in the range [0,1] with spacing 0.05 and we also chose σ = 0.05. This resulted in a total of *N_Q_* = 21 umbrella potentials.

Each set of isolated domain simulations was performed using 8 walkers, with *T_max_* = 480 K and *T_min_* = 300 K. Further details can be found in our previous work,^81^ where we also demonstrated that MM-OPES simulations provide the same accuracy as PTWTM-WTM simulations under optimal conditions. Thus, the results from the simulations are comparable.

After initial solvation, systems were equilibrated with 100 ps NVT simulations followed by 100 ps NPT (P=1 bar) simulations for three temperatures. The first equilibration set of NVT and then NPT simulations was done at room temperature (T=300 K), then at the highest temperature (T=440 K for full-length simulations and T=480 K for isolated domain simulations), and then at their run temperatures prior to starting the simulations. It should be noted that the equilibration at the highest temperature made us sure that the system would not collapse during the course of simulations.

The run time for simulations per replica is as below: 1.7 µs for each main full-length simulation replica, 1.5 µs for each restricted-folded simulation replica, and 1.1 µs for each isolated domain simulation walker. Therefore, these simulation sets yielded the aggregated computational time of 34 µs for the main full-length simulation, 30 µs for the restricted-folded simulations, and 8.8 µs for each isolated domain simulation.

All simulations were performed using the GROMACS code (version 2021.4) with the PLUMED (version 2.8) patch for sampling.

### Convergence

To assess the convergence of our simulations, we conducted a comprehensive analysis, summarized in Extended Data Fig. 2-4. We monitored several structural observables as a function of time, including the global and domain-level parameters such as R_g_, the collective variable Q for the full-length SpA and Q for individual domains. For each observable, we plotted time series data and overlaid 50 ns running averages to identify equilibration behavior (Extended Data Fig. 2-3). In all cases, a conservative equilibration window was discarded, and subsequent plateaus or stable fluctuations were taken as evidence of convergence.

As the most stringent test for convergence, we calculated the free energy difference between the folded and unfolded states (ΔG = —k_B_T ln(F_F_ /F_U_)) as a function of simulation time (Extended Data Fig. 4). Folded and unfolded populations (F_F_and F_U_) were estimated using Boltzmann-weighted sampling within respective basins in Q space (folded: Q > 0.8; unfolded: Q < 0.2). We assumed convergence when ΔG stabilized within *±*k_B_T (∼2.5 kJ/mol) of its final value, indicating that transitions between folded and unfolded states were adequately sampled and that the computed thermodynamics reflect equilibrium behavior within the range of thermal fluctuations.

Together, these analyses confirm that all production runs used in this study are sufficiently converged for both thermodynamic and structural interpretation.

### Simulations Analyses

We projected free energy onto different order parameters: *Q*, radius of gyration (*R_g_*), or dihedral angles (*ϕ*and *ψ*). Among these parameters, Q is the collective variable biased during the course of simulations, besides potential energy. When Q includes atom pairs of all domains and domain X, is denoted as simply Q, but when Q is calculated based on atom pairs within a single domain, it is specified as *Q_E_, Q_D_, Q_A_, Q_B_*, or *Q_C_*.

All free energy plots presented are unbiased, meaning that after reweighting all bias deposited on Q and potential energy, the resulting unbiased probability densities were used to calculate free energy. We discarded the initial 600 ns/replica of full-length simulations and the initial (> 500 ns/replica of isolated domains simulations as equilibration time, and the remaining data (> 500 ns/replica) for analysis.

To confirm equilibration, we examined Q and *R_g_* as a function of time (Extended Data Figs. 2-3) to assess whether they fluctate around an average (i.e., no drift). We also calculated the free energy difference between unfolded and folded states, denoted as Δ*G*, as a function of time (Extended Data Fig. 4). Equilibration was defined as the point where Δ*G* remained stable within thermal fluctuations (*±*kT ≈ 2.5 kJ/mol at 300 K). This criterion aligns well with the low fluctuations regime in the running average of Q and R_g_ over sampling time (Extended Data Figs. 2-3).

To estimate HC from simulations, the Dictionary of Secondary Structures of Proteins (DSSP) algorithm^30^ on a per-residue basis was used.

Calculations of *R_g_* were done based on heavy backbone atoms. Internal distances were computed every nanosecond from the simulation trajectories, and unbiased ensemble-averaged internal distances < *R_ij_* > were obtained using weighted histograms.

### Protein source and preparation

Recombinant SpA (rSpA) 50 mg/ml in purified water was obtained from Repligen. It was diluted and buffer-exchanged into PBS using a Zeba Spin Desalting column (0.5 mL, Thermo Fisher Scientific) with a molecular weight cutoff (MWCO) of 40 kDa for rSpA and 7 kDa for the Z1 construct.

Domain Z was expressed, purified, and characterized largely as described in our previous work.^84^ It was expressed using a vector provided in BL21 DE3 *E. coli*. Purification was performed using Ni-NTA + SEC, and validation of molecular mass was done using SDS-PAGE and MALDI, in accordance with the protocol described in a previous paper.

### Circular dichroism spectroscopy (CD)

We determined helix fractions from CD data as described by Zavrtanik.^85^ CD was performed on an OLIS RSM 1000 CD with a Xenon lamp. A Julabo F30-C heated-refrigerated circulating water bath was used for temperature control. The monochromator was calibrated using *(1S)-(+)-camphor-10-sulfonic acid* (also known as the CSA standard^86^), a standard compound with a characteristic ellipticity band at 291 nm used to verify instrument calibration in CD spectroscopy. All samples were generated using a 2 mg/ml protein concentration in PBS buffer. Before each run, air and water blanks were performed to verify the cleanliness of the cuvettes, which were 1 mm path length. The buffer blank was applied as the baseline. Measurements were performed in one-nanometer steps over a wavelength range of 330 nm to 200 nm. The temperature was scanned from 20 *^○^*C to 90 *^○^*C in increments of 10 *^○^*C, with 5 minutes of stabilization at each temperature, a time confirmed by a probe on the sample chamber to be sufficient for temperature stabilization.

### Differential Scanning Calorimetry (DSC)

DSC samples were prepared in 10 mM phosphate buffer and 50 mM NaCl at pH 7 with the sample concentration of 100 µM. Samples were sent to the Huck Institute at Penn State and run on a MicroCal VP-Capillary DSC with a Tantalum 130 µl capillary type cell. Samples were diluted to 1 mg/ml and run at 1 °C/ min from 15 to 130 °C. Scans were run in triplicate for each sample provided. Bovine serum albumin (BSA) protein was used as a control.

## Acknowledgements

We thank Suman Nandy, PhD, and Vijay Maranholkar, PhD, for helpful discussions. GHZ is a Cancer Prevention and Research Institute of Texas (CPRIT) scholar in cancer research and is supported by CPRIT-RR220008, the Welch Foundation (Award E-2221, and Catalyst Center for Advanced Bioactive Materials Crystallization Award V-E-0001), and National Science Foundation CBET-2442006 (CAREER). RCW is supported by the National Institutes of Health (1R61AI174294). The simulations presented in this work were performed on the computational resources provided by the Hewlett-Packard Enterprise Data Science Institute at the University of Houston.

## References

(1) Qi, Y.; Martin, J. W.; Barb, A. W.; Thélot, F.; Yan, A. K.; Donald, B. R.; Oas, T. G. Continuous interdomain orientation distributions reveal components of binding thermodynamics. Journal of molecular biology 2018, 430, 3412–3426.

(2) Capp, J. A.; Hagarman, A.; Richardson, D. C.; Oas, T. G. The statistical conformation of a highly flexible protein: small-angle X-ray scattering of S. aureus protein A. Structure 2014, 22, 1184–1195.

(3) Pagano, L.; Malagrinò, F.; Visconti, L.; Troilo, F.; Pennacchietti, V.; Nardella, C.; Toto, A.; Gianni, S. Probing the effects of local frustration in the folding of a multidomain protein. Journal of Molecular Biology 2021, 433, 167087.

(4) Deis, L. N.; Pemble, C. W.; Qi, Y.; Hagarman, A.; Richardson, D. C.; Richardson, J. S.; Oas, T. G. Multiscale conformational heterogeneity in staphylococcal protein a: possible determinant of functional plasticity. Structure 2014, 22, 1467–1477.

(5) Cui, X.; Ge, L.; Chen, X.; Lv, Z.; Wang, S.; Zhou, X.; Zhang, G. Beyond static structures: protein dynamic conformations modeling in the post-AlphaFold era. Briefings in bioinformatics 2025, 26, bbaf340.

(6) Lin, R. L. F.; Bellaiche, A.; Etchebest, C. The key role of the dynamics and flexibility of proteins in functional mechanisms: How computational methods can contribute to their identification. Biochimie 2025,

(7) Lesovoy, D.; Roshchin, K.; Sala, B. M.; Sandalova, T.; Achour, A.; Agback, T.; Agback, P.; Orekhov, V. Accurate Protein Dynamic Conformational Ensembles: Combining AlphaFold, MD, and Amide 15N (1H) NMR Relaxation. International Journal of Molecular Sciences 2025, 26, 8917.

(8) Gershenson, A.; Gosavi, S.; Faccioli, P.; Wintrode, P. L. Successes and challenges in simulating the folding of large proteins. Journal of Biological Chemistry 2020, 295, 15–33.

(9) Grasso, D.; Galderisi, S.; Santucci, A.; Bernini, A. Pharmacological chaperones and protein conformational diseases: approaches of computational structural biology. International Journal of Molecular Sciences 2023, 24, 5819.

(10) Jackson, S. E. How do small single-domain proteins fold? Folding and Design 1998, 3, R81–R91.

(11) Zhuravlev, P. I.; Hinczewski, M.; Chakrabarti, S.; Marqusee, S.; Thirumalai, D. Force-dependent switch in protein unfolding pathways and transition-state movements. Proceedings of the National Academy of Sciences 2016, 113, E715–E724.

(12) Schuler, B.; Lipman, E. A.; Eaton, W. A. Probing the free-energy surface for protein folding with single-molecule fluorescence spectroscopy. Nature 2002, 419, 743–747.

(13) Onuchic, J. N.; Wolynes, P. G. Theory of protein folding. Current opinion in structural biology 2004, 14, 70–75.

(14) Han, J.-H.; Batey, S.; Nickson, A. A.; Teichmann, S. A.; Clarke, J. The folding and evolution of multidomain proteins. Nature reviews Molecular cell biology 2007, 8, 319– 330.

(15) Borgia, M. B.; Borgia, A.; Best, R. B.; Steward, A.; Nettels, D.; Wunderlich, B.; Schuler, B.; Clarke, J. Single-molecule fluorescence reveals sequence-specific misfolding in multidomain proteins. Nature 2011, 474, 662–665.

(16) Laursen, L.; Gianni, S.; Jemth, P. Dissecting inter-domain cooperativity in the folding of a multi domain protein. Journal of Molecular Biology 2021, 433, 167148.

(17) Santorelli, D.; Marcocci, L.; Pennacchietti, V.; Nardella, C.; Diop, A.; Pietrangeli, P.; Pagano, L.; Toto, A.; Malagrinò, F.; Gianni, S. Understanding the molecular basis of folding cooperativity through a comparative analysis of a multidomain protein and its isolated domains. Journal of Biological Chemistry 2023, 299.

(18) Vergara, R.; Berrocal, T.; Juárez Mejía, E. I.; Romero-Romero, S.; Velázquez-López, I.; Pulido, N. O.; López Sanchez, H. A.; Silva, D.-A.; Costas, M.; Rodríguez-Romero, A.; others Thermodynamic and kinetic analysis of the LAO binding protein and its isolated domains reveal non-additivity in stability, folding and function. The FEBS journal 2023, 290, 4496–4512.

(19) Braselmann, E.; Chaney, J. L.; Clark, P. L. Folding the proteome. Trends in biochemical sciences 2013, 38, 337–344.

(20) Falugi, F.; Kim, H. K.; Missiakas, D. M.; Schneewind, O. Role of protein A in the evasion of host adaptive immune responses by Staphylococcus aureus. MBio 2013, 4, 10–1128.

(21) Graille, M.; Stura, E. A.; Corper, A. L.; Sutton, B. J.; Taussig, M. J.; Charbonnier, J.-B.; Silverman, G. J. Crystal structure of a Staphylococcus aureus protein A domain complexed with the Fab fragment of a human IgM antibody: structural basis for recognition of B-cell receptors and superantigen activity. Proceedings of the National Academy of Sciences 2000, 97, 5399–5404.

(22) Palmqvist, N.; Silverman, G. J.; Josefsson, E.; Tarkowski, A. Bacterial cell wall-expressed protein A triggers supraclonal B-cell responses upon in vivo infection with Staphylococcus aureus. Microbes and infection 2005, 7, 1501–1511.

(23) Gómez, M. I.; Lee, A.; Reddy, B.; Muir, A.; Soong, G.; Pitt, A.; Cheung, A.; Prince, A. Staphylococcus aureus protein A induces airway epithelial inflammatory responses by activating TNFR1. Nature medicine 2004, 10, 842–848.

(24) O’Seaghdha, M.; van Schooten, C. J.; Kerrigan, S. W.; Emsley, J.; Silverman, G. J.; Cox, D.; Lenting, P. J.; Foster, T. J. Staphylococcus aureus protein A binding to von Willebrand factor A1 domain is mediated by conserved IgG binding regions. The FEBS journal 2006, 273, 4831–4841.

(25) Thammavongsa, V.; Kim, H. K.; Missiakas, D.; Schneewind, O. Staphylococcal manipulation of host immune responses. Nature Reviews Microbiology 2015, 13, 529–543.

(26) Deis, L. N.; Wu, Q.; Wang, Y.; Qi, Y.; Daniels, K. G.; Zhou, P.; Oas, T. G. Suppression of conformational heterogeneity at a protein–protein interface. Proceedings of the National Academy of Sciences 2015, 112, 9028–9033.

(27) Rigi, G.; Ghaedmohammadi, S.; Ahmadian, G. A comprehensive review on staphylococcal protein A (SpA): Its production and applications. Biotechnology and applied biochemistry 2019, 66, 454–464.

(28) Cruz, A. R.; Boer, M. A. d.; Strasser, J.; Zwarthoff, S. A.; Beurskens, F. J.; de Haas, C. J.; Aerts, P. C.; Wang, G.; de Jong, R. N.; Bagnoli, F.; others Staphylococcal protein A inhibits complement activation by interfering with IgG hexamer formation. Proceedings of the National Academy of Sciences 2021, 118, e2016772118.

(29) Romih, T.; Konjevíc, I.; Žibret, L.; Fazarinc, I.; Beltram, A.; Majer, D.; Finšgar, M.; Hočevar, S. B. The effect of preconditioning strategies on the adsorption of model proteins onto screen-printed carbon electrodes. Sensors 2022, 22, 4186.

(30) Kabsch, W.; Sander, C. Dictionary of protein secondary structure: pattern recognition of hydrogen-bonded and geometrical features. Biopolymers: Original Research on Biomolecules 1983, 22, 2577–2637.

(31) Haley, O. C.; Tibbs-Cortes, L.; Hayford, R. K.; Harding, S.; Woodhouse, M.; Cannon, E.; Gardiner, J.; Portwood III, J.; Sen, T. Z.; Kim, H.-S.; others Why do some predicted protein structures fold poorly? Benchmarking AlphaFold, ESMFold, and Boltz in maize. bioRxiv 2025, 2025–07.

(32) Jumper, J.; Evans, R.; Pritzel, A.; Green, T.; Figurnov, M.; Ronneberger, O.; Tunyasuvunakool, K.; Bates, R.; Žídek, A.; Potapenko, A.; others Highly accurate protein structure prediction with AlphaFold. Nature 2021, 596, 583–589.

(33) Baek, M.; DiMaio, F.; Anishchenko, I.; Dauparas, J.; Ovchinnikov, S.; Lee, G. R.; Wang, J.; Cong, Q.; Kinch, L. N.; Schaeffer, R. D.; others Accurate prediction of protein structures and interactions using a three-track neural network. Science 2021, 373, 871–876.

(34) Lin, Z.; Akin, H.; Rao, R.; Hie, B.; Zhu, Z.; Lu, W.; Smetanin, N.; Verkuil, R.; Kabeli, O.; Shmueli, Y.; others Evolutionary-scale prediction of atomic-level protein structure with a language model. Science 2023, 379, 1123–1130.

(35) Passaro, S.; Corso, G.; Wohlwend, J.; Reveiz, M.; Thaler, S.; Somnath, V. R.; Getz, N.; Portnoi, T.; Roy, J.; Stark, H.; others Boltz-2: Towards accurate and efficient binding affinity prediction. BioRxiv 2025, 2025–06.

(36) Emenecker, R. J.; Griffith, D.; Holehouse, A. S. Metapredict: a fast, accurate, and easy-to-use predictor of consensus disorder and structure. Biophysical journal 2021, 120, 4312–4319.

(37) Uversky, V. N.; Gillespie, J. R.; Fink, A. L. Why are “natively unfolded” proteins unstructured under physiologic conditions? Proteins: structure, function, and bioinformatics 2000, 41, 415–427.

(38) Starovasnik, M. A.; O’Connell, M. P.; Fairbrother, W. J.; Kelley, R. F. Antibody variable region binding by Staphylococcal protein A: thermodynamic analysis and location of the Fv binding site on E-domain. Protein Science 1999, 8, 1423–1431.

(39) Shea, J.-E.; Onuchic, J. N.; Brooks III, C. L. Exploring the origins of topological frustration: design of a minimally frustrated model of fragment B of protein A. Proceedings of the National Academy of Sciences 1999, 96, 12512–12517.

(40) Myrhammar, A.; Rosik, D.; Karlstrom, A. E. Photocontrolled reversible binding between the protein A-derived Z domain and immunoglobulin G. Bioconjugate chemistry 2020, 31, 622–630.

(41) Alonso, D. O.; Daggett, V. Staphylococcal protein A: unfolding pathways, unfolded states, and differences between the B and E domains. Proceedings of the National Academy of Sciences 2000, 97, 133–138.

(42) Cedergren, L.; Andersson, R.; Jansson, B.; Uhlén, M.; Nilsson, B. Mutational analysis of the interaction between staphylococcal protein A and human IgG1. *Protein Engineering*, Design and Selection 1993, 6, 441–448.

(43) Zerze, G. H.; Uz, B.; Mittal, J. Folding thermodynamics of β-hairpins studied by replica-exchange molecular dynamics simulations. *Proteins: Structure*, Function, and Bioinformatics 2015, 83, 1307–1315.

(44) Du, D.; Gai, F. Understanding the folding mechanism of an α-helical hairpin. Biochemistry 2006, 45, 13131–13139.

(45) Yang, J. S.; Wallin, S.; Shakhnovich, E. I. Universality and diversity of folding mechanics for three-helix bundle proteins. Proceedings of the National Academy of Sciences 2008, 105, 895–900.

(46) Sato, S.; Religa, T. L.; Daggett, V.; Fersht, A. R. Testing protein-folding simulations by experiment: B domain of protein A. Proceedings of the National Academy of Sciences 2004, 101, 6952–6956.

(47) Pereira, A. F.; Martinez, L. Helical Content Correlations and Hydration Structures of the Folding Ensemble of the B Domain of Protein A. Journal of Chemical Information and Modeling 2024, 64, 3350–3359.

(48) Myers, J. K.; Oas, T. G. Preorganized secondary structure as an important determinant of fast protein folding. Nature structural biology 2001, 8, 552–558.

(49) Flory, P. J. Principles of polymer chemistry; Cornell university press, 1953.

(50) Zerze, G. H.; Best, R. B.; Mittal, J. Sequence-and temperature-dependent properties of unfolded and disordered proteins from atomistic simulations. The Journal of Physical Chemistry B 2015, 119, 14622–14630.

(51) Alston, J. J.; Ginell, G. M.; Soranno, A.; Holehouse, A. S. The analytical Flory random coil is a simple-to-use reference model for unfolded and disordered proteins. The Journal of Physical Chemistry B 2023, 127, 4746–4760.

(52) Rubinstein, M.; Colby, R. H. Polymer physics; Oxford university press, 2003.

(53) Das, R. K.; Pappu, R. V. Conformations of intrinsically disordered proteins are influenced by linear sequence distributions of oppositely charged residues. Proceedings of the National Academy of Sciences 2013, 110, 13392–13397.

(54) Rajasekaran, N.; Kaiser, C. M. Navigating the complexities of multi-domain protein folding. Current Opinion in Structural Biology 2024, 86, 102790.

(55) Mazigi, O.; Schofield, P.; Langley, D. B.; Christ, D. Protein A superantigen: structure, engineering and molecular basis of antibody recognition. *Protein Engineering*, Design and Selection 2019, 32, 359–366.

(56) Galpern, E. A.; Marchi, J.; Mora, T.; Walczak, A. M.; Ferreiro, D. U. Evolution and folding of repeat proteins. Proceedings of the National Academy of Sciences 2022, 119, e2204131119.

(57) Bhaskara, R. M.; Srinivasan, N. Stability of domain structures in multi-domain proteins. Scientific reports 2011, 1, 40.

(58) Bowman, G. R. Alphafold and protein folding: Not dead yet! The frontier is conformational ensembles. Annual review of biomedical data science 2024, 7, 51–57.

(59) Niazi, S. K.; Yang, J. A comprehensive application of FiveFold for conformation ensemble-based protein structure prediction. Scientific Reports 2025, 15, 33498.

(60) Kalakoti, Y.; Wallner, B. AFsample2 predicts multiple conformations and ensembles with AlphaFold2. Communications Biology 2025, 8, 373.

(61) Monteiro da Silva, G.; Cui, J. Y.; Dalgarno, D. C.; Lisi, G. P.; Rubenstein, B. M. High-throughput prediction of protein conformational distributions with subsampled AlphaFold2. Nature Communications 2024, 15, 2464.

(62) Chakravarty, D.; Lee, M.; Porter, L. Proteins with alternative folds reveal blind spots in AlphaFold-based protein structure prediction. arXiv. arXiv preprint arXiv: 2410.14898 2024,

(63) McDonald, E. F.; Jones, T.; Plate, L.; Meiler, J.; Gulsevin, A. Benchmarking AlphaFold2 on peptide structure prediction. Structure 2023, 31, 111–119.

(64) Agarwal, V.; McShan, A. C. The power and pitfalls of AlphaFold2 for structure prediction beyond rigid globular proteins. Nature chemical biology 2024, 20, 950–959.

(65) Schneewind, O.; Missiakas, D. M. Staphylococcal protein secretion and envelope assembly. Microbiology spectrum 2019, 7, 10–1128.

(66) Makepeace, K. A.; Brodie, N. I.; Popov, K. I.; Gudavicius, G.; Nelson, C. J.; Petrotchenko, E. V.; Dokholyan, N. V.; Borchers, C. H. Ligand-induced disorder-to-order transitions characterized by structural proteomics and molecular dynamics simulations. Journal of proteomics 2020, 211, 103544.

(67) Fujino, Y.; Miyagawa, T.; Torii, M.; Inoue, M.; Fujii, Y.; Okanishi, H.; Kanai, Y.; Masui, R. Structural changes induced by ligand binding drastically increase the thermostability of the Ser/Thr protein kinase TpkD from Thermus thermophilus HB8. FEBS letters 2021, 595, 264–274.

(68) Stenström, O.; Diehl, C.; Modig, K.; Akke, M. Ligand-induced protein transition state stabilization switches the binding pathway from conformational selection to induced fit. Proceedings of the National Academy of Sciences 2024, 121, e2317747121.

(69) Lewis, S.; Hempel, T.; Jiménez-Luna, J.; Gastegger, M.; Xie, Y.; Foong, A. Y.; Satorras, V. G.; Abdin, O.; Veeling, B. S.; Zaporozhets, I.; others Scalable emulation of protein equilibrium ensembles with generative deep learning. Science 2025, eadv9817.

(70) Huang, J.; Rauscher, S.; Nawrocki, G.; Ran, T.; Feig, M.; De Groot, B. L.; Grubmüller, H.; MacKerell Jr, A. D. CHARMM36m: an improved force field for folded and intrinsically disordered proteins. Nature methods 2017, 14, 71–73.

(71) Barducci, A.; Bussi, G.; Parrinello, M. Well-tempered metadynamics: a smoothly converging and tunable free-energy method. Physical review letters 2008, 100, 020603.

(72) Bonomi, M.; Parrinello, M. Enhanced sampling in the well-tempered ensemble. Physical review letters 2010, 104, 190601.

(73) Bussi, G.; Gervasio, F. L.; Laio, A.; Parrinello, M. Free-energy landscape for β hairpin folding from combined parallel tempering and metadynamics. Journal of the American Chemical Society 2006, 128, 13435–13441.

(74) Gil-Ley, A.; Bussi, G. Enhanced conformational sampling using replica exchange with collective-variable tempering. Journal of chemical theory and computation 2015, 11, 1077–1085.

(75) Best, R. B.; Mittal, J. Balance between α and β structures in ab initio protein folding. The Journal of Physical Chemistry B 2010, 114, 8790–8798.

(76) Best, R. B.; Hummer, G.; Eaton, W. A. Native contacts determine protein folding mechanisms in atomistic simulations. Proc. Nat. Acad. Sci. 2013, 110, 17874–17879.

(77) Zerze, G. H.; Stillinger, F. H.; Debenedetti, P. G. Thermodynamics of DNA Hybridization from Atomistic Simulations. J. Phys. Chem. B 2021, 125, 771–779.

(78) Zerze, G. H.; Piaggi, P. M.; Debenedetti, P. G. A Computational Study of RNA Tetraloop Thermodynamics, Including Misfolded States. J. Phys. Chem. B 2021, 125, 13685–13695.

(79) McGovern, M.; De Pablo, J. A boundary correction algorithm for metadynamics in multiple dimensions. The Journal of chemical physics 2013, 139.

(80) Invernizzi, M.; Piaggi, P. M.; Parrinello, M. Unified approach to enhanced sampling. Phys. Rev. X 2020, 10, 041034.

(81) Rahimi, K.; Piaggi, P. M.; Zerze, G. H. Comparison of on-the-fly probability enhanced sampling and parallel tempering combined with metadynamics for atomistic simulations of RNA tetraloop folding. The Journal of Physical Chemistry B 2023, 127, 4722–4732.

(82) Piaggi, P. M.; Parrinello, M. Calculation of phase diagrams in the multithermal-multibaric ensemble. J. Chem. Phys. 2019, 150, 244119.

(83) Malekzadeh, K.; Zerze, G. H. Optimizing On-the-Fly Probability Enhanced Sampling for Complex RNA Systems: Sampling Free Energy Surfaces of an H-Type Pseudoknot. Journal of Chemical Information and Modeling 2025, 65, 3605–3614.

(84) Nandy, S.; Maranholkar, V. M.; Crum, M.; Wasden, K.; Patil, U.; Goyal, A.; Vu, B.; Kourentzi, K.; Mo, W.; Henrickson, A.; others Expression and characterization of intein-cyclized trimer of Staphylococcus aureus protein A domain Z. International journal of molecular sciences 2023, 24, 1281.

(85) Zavrtanik, U.; Lah, J.; Hadži, S. Estimation of peptide helicity from circular dichroism using the ensemble model. J. Phys. Chem. B 2024, 128, 2652–2663.

(86) Greenfield, N. J. Using circular dichroism spectra to estimate protein secondary structure. Nat. Protoc. 2006, 1, 2876–2890.

