## Supporting Figures and Tables for "Full-Length Context Disrupts Folding of IgG-Binding Domains of Protein A"

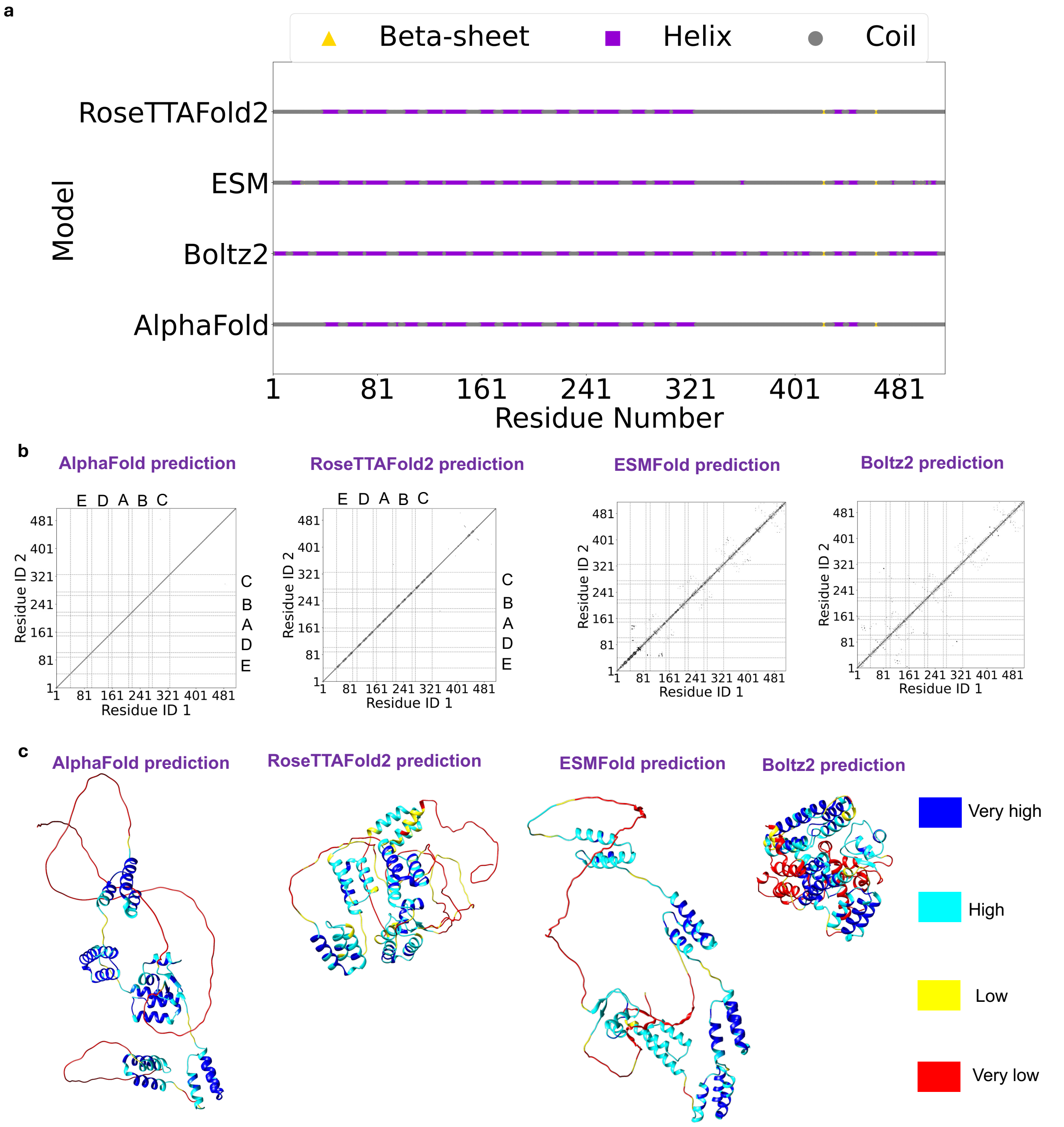


**Extended Data Fig. 1** | **a,** DSSP analysis for the SpA structures predicted from different AI models. **b,** heavy atoms contact maps for four AI models using a cutoff of 2.5 Å. Note that most of these contacts that form within (less than) 2.5 Å distance are unphysical steric clashes. **c,** AI-predicted structures colored by confidence.

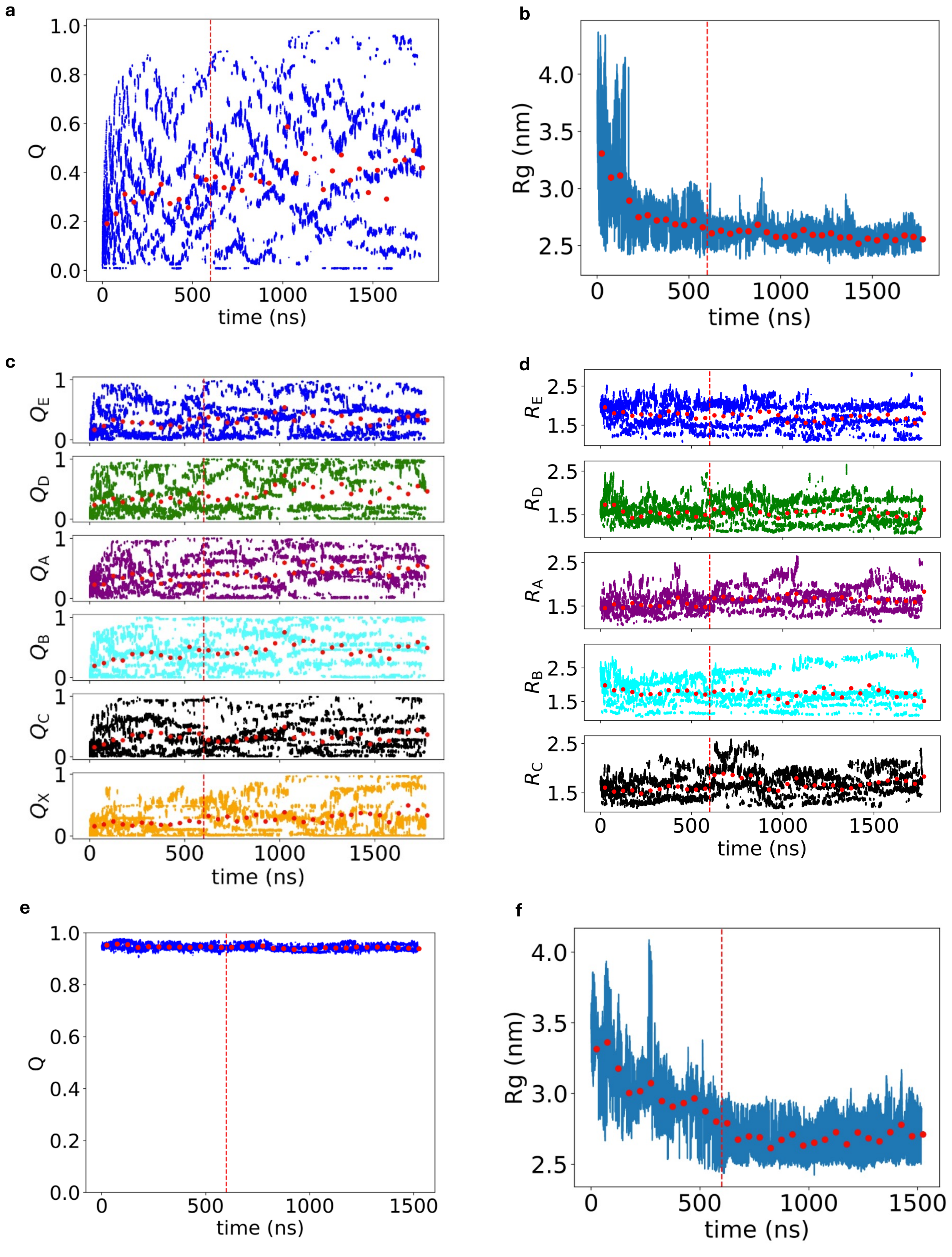


**Extended Data Fig. 2** | Time evolution of the collective variable Q and $R_g$ at 300 K for (**a,b**) main full-length simulation, (**c,d**) for each domain of SpA in the main full-length simulation, (**e,f**) for the restricted-folded full-length simulation. Data up to the red vertical line is discarded as equilibration, and the dashed red line represents the running average (with 50 ns blocks).

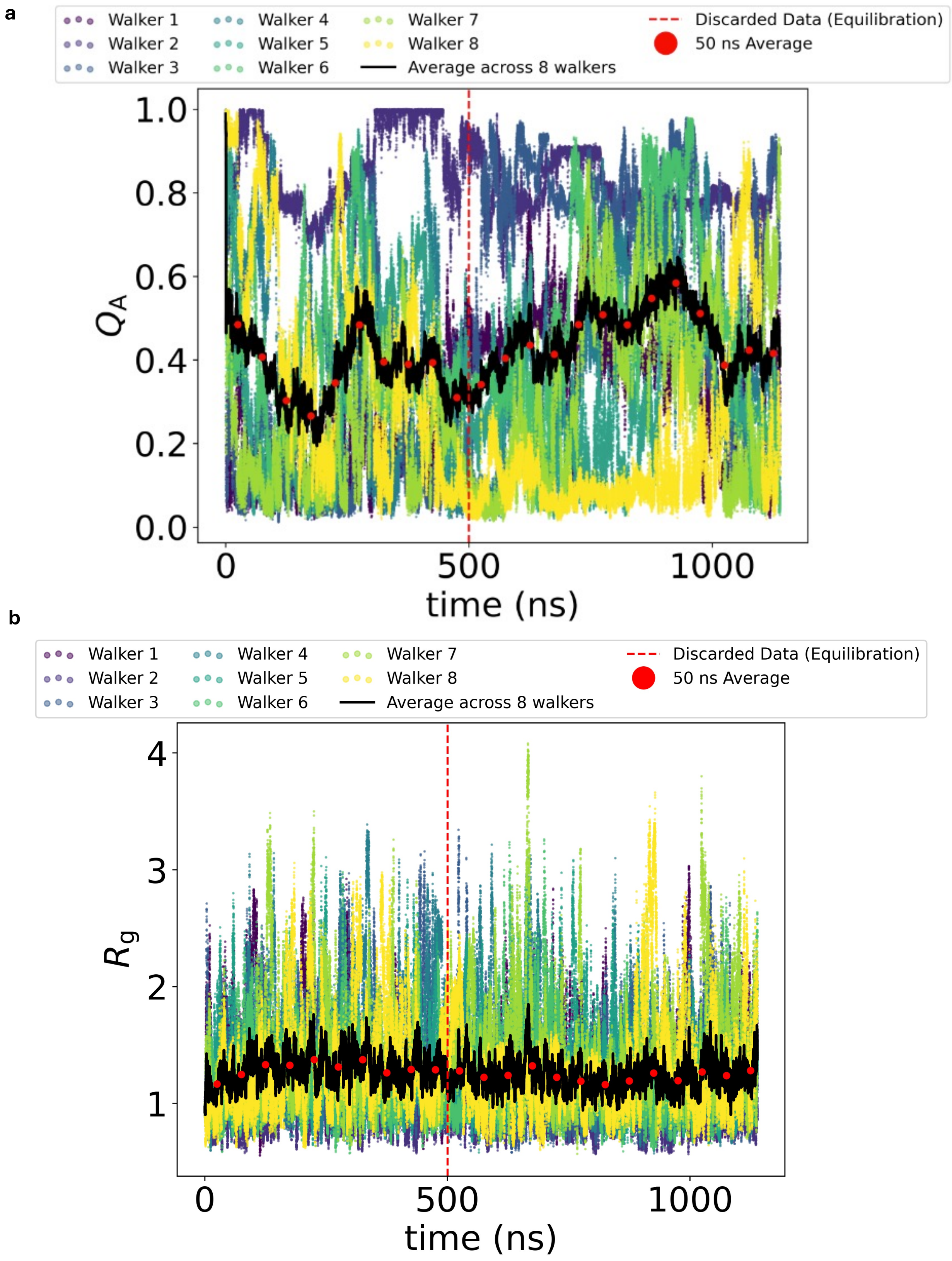


**Extended Data Fig. 3** | **a,** Time evolution of collective variable Q for each walker of the MM-OPES simulations, for isolated domain A. **b,** Time evolution of $R_g$ for each walker for the isolated domain A simulations. Data up to the red vertical line is discarded as equilibration, and the red dots represent a running average (for 50 ns/replica windows).

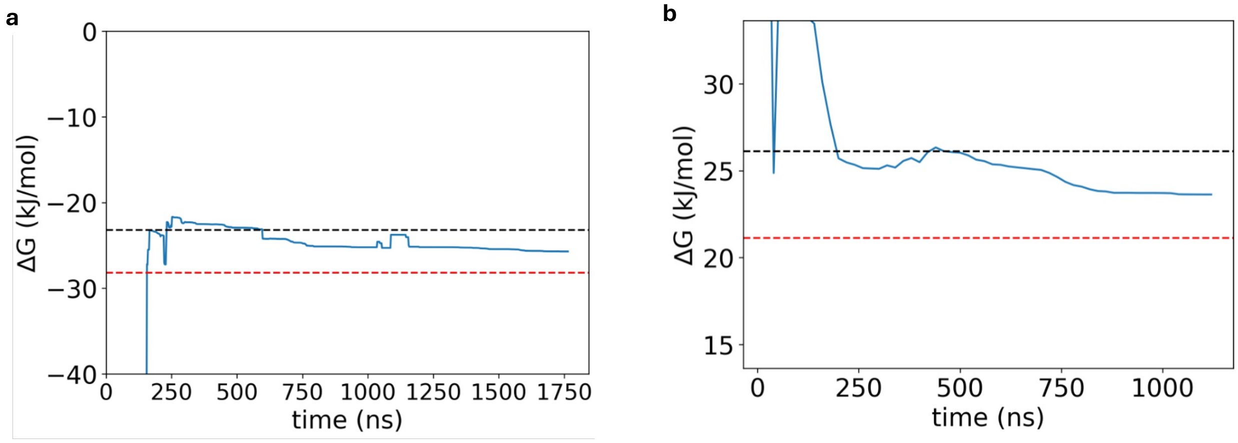


**Extended Data Fig. 4** | The change in free energy difference between the folded (F) and unfolded (U) states as a function of time ($\Delta G = -kT \ln \frac{F_F}{F_U}$ ). $F_U$ is calculated as the Boltzman average of the unfolded basin $Q < 0.2$, and $F_F$ is calculated as the Boltzman average of the folded basin $Q > 0.8$. The dashed lines show $\Delta G \pm kT$ ($\approx 2.5$ kJ/mol) from the last frame. **a,** for the main simulation at 300 K **b,** for the isolated domain A simulations.

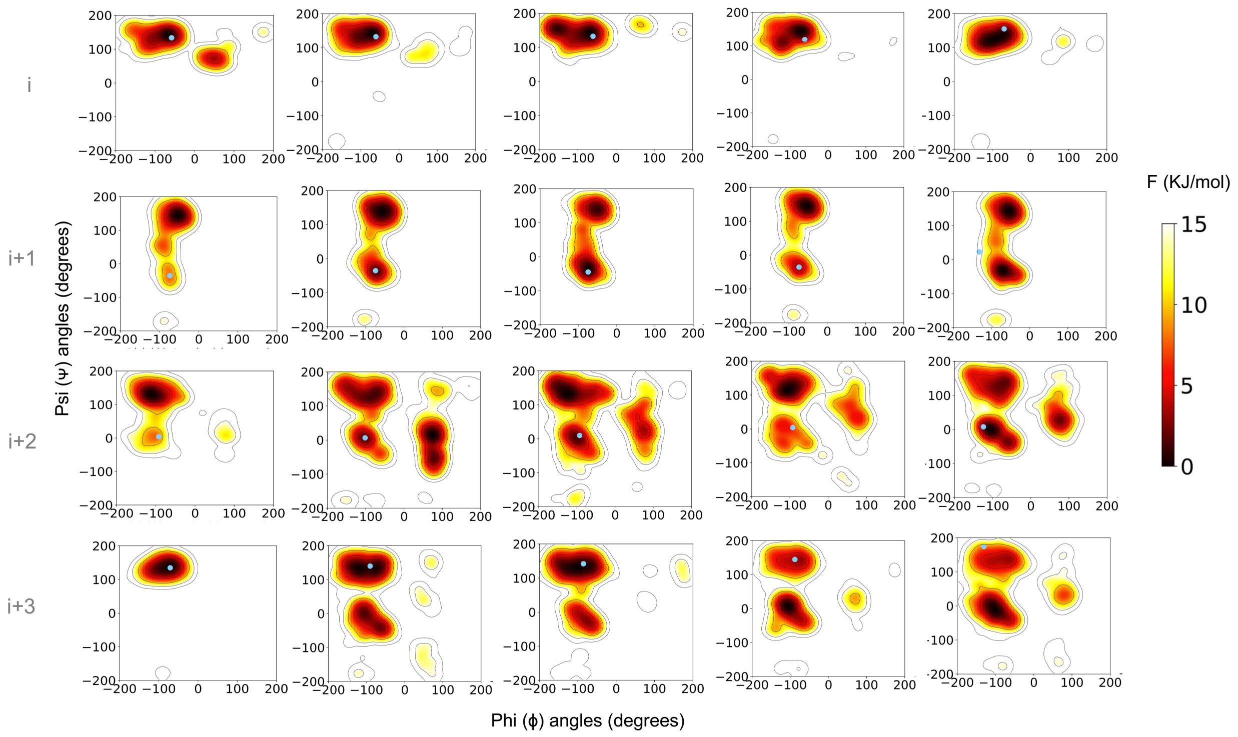


**Extended Data Fig. 5** | Free energy surface projected onto the dihedral angles for the first turn residues in the triple-$\alpha$-helical structure of individual domains of SpA when they are in the full-length assembly. The domains are ordered E, D, A, B, and C from left to right.

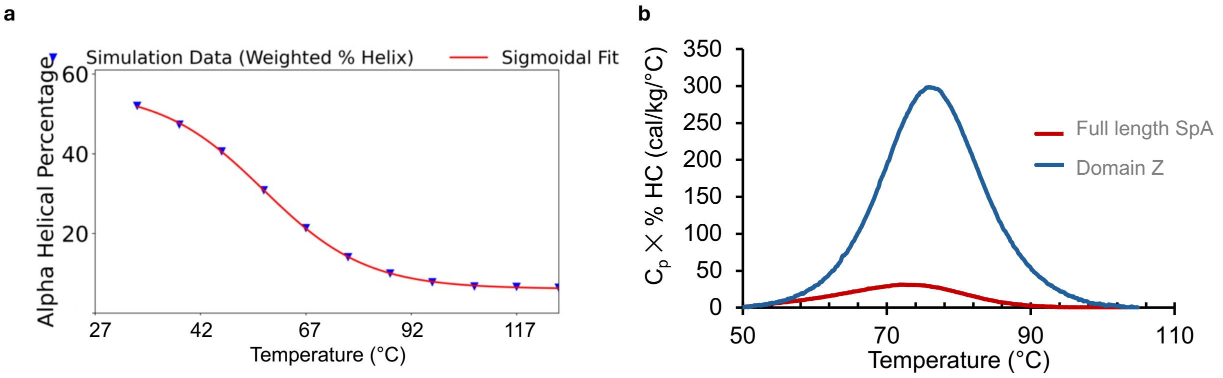


**Extended Data Fig. 6** | **a,** Helix percentage of domain Z as a function of temperature calculated from weighted DSSP **b,** Helix-content–weighted heat capacity (Cp × HC) profiles for full-length SpA and domain Z.

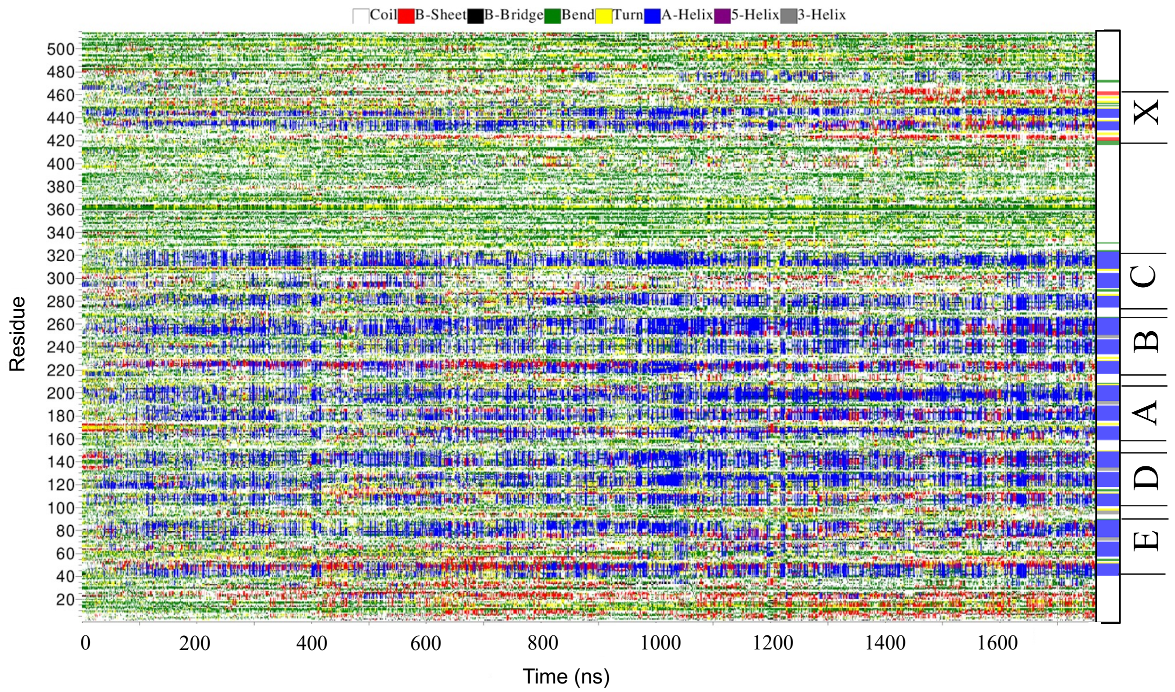


**Extended Data Fig. 7** | Color-coded DSSP analysis (not reweighted) as a function of simulation time for the main full-length simulation at 300 K. The rightmost bar displays the DSSP analysis of the AlphaFold-predicted structure, with domains E, D, A, B, C, and domain X labeled for reference.

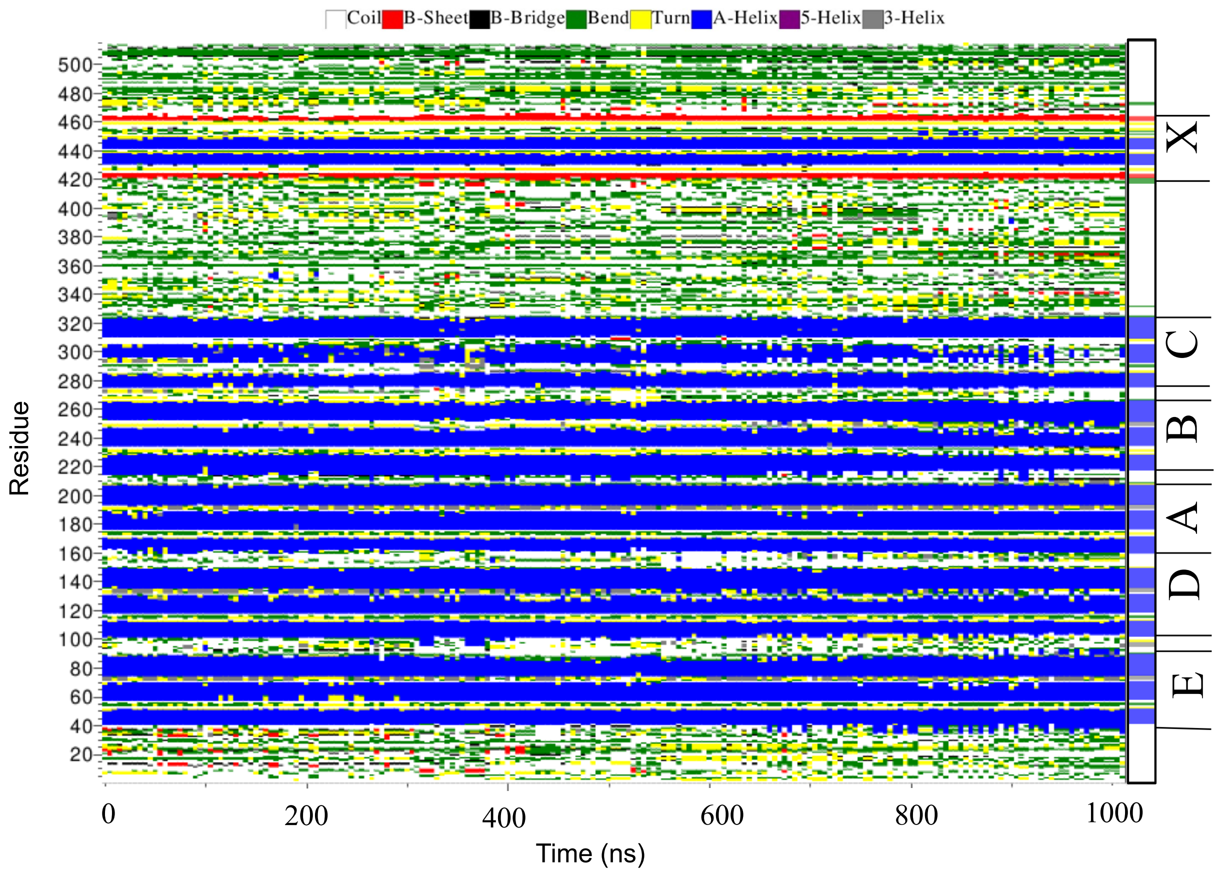


**Extended Data Fig. 8** | Color-coded DSSP analysis (not reweighted) as a function of simulation time for the restricted-folded full-length simulation at 300 K. The rightmost bar displays the DSSP analysis of the AlphaFold-predicted structure, with domains E, D, A, B, C, and domain X labeled for reference.

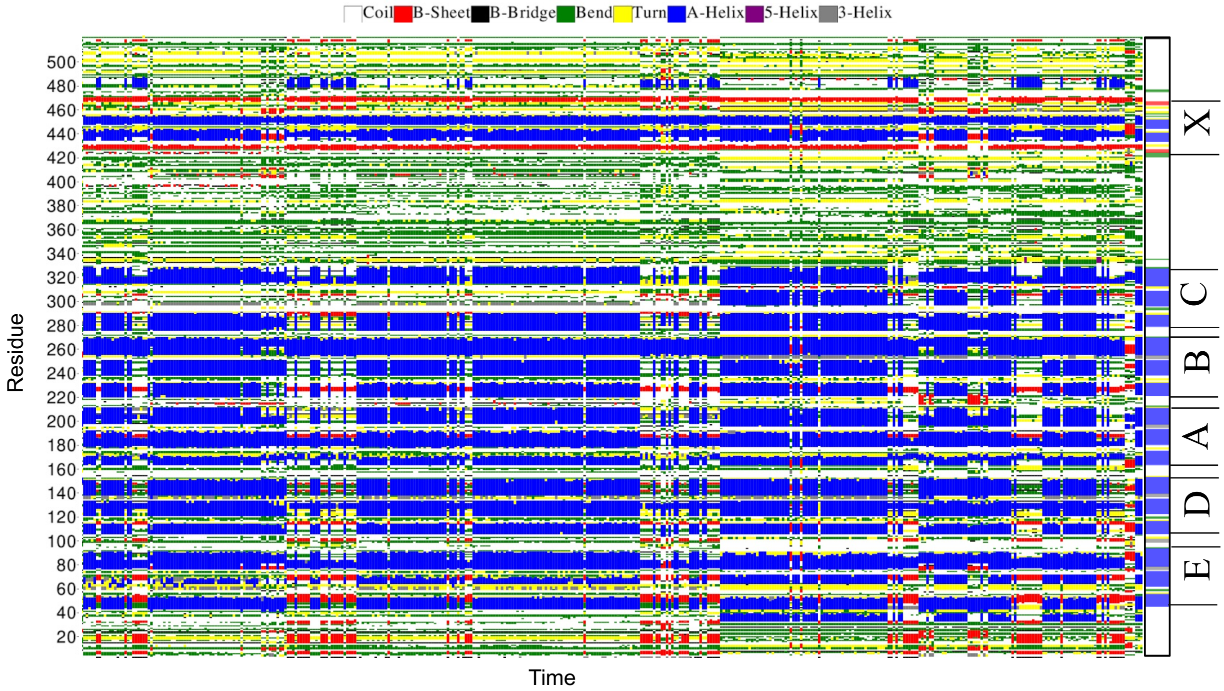


**Extended Data Fig. 9** | Color-coded DSSP analysis (not reweighted) as a function of simulation time for the subtrajectory $Q > 0.9$ of the main simulation. The rightmost bar displays the DSSP analysis of the AlphaFold-predicted structure, with domains E, D, A, B, C, and domain X labeled for reference.

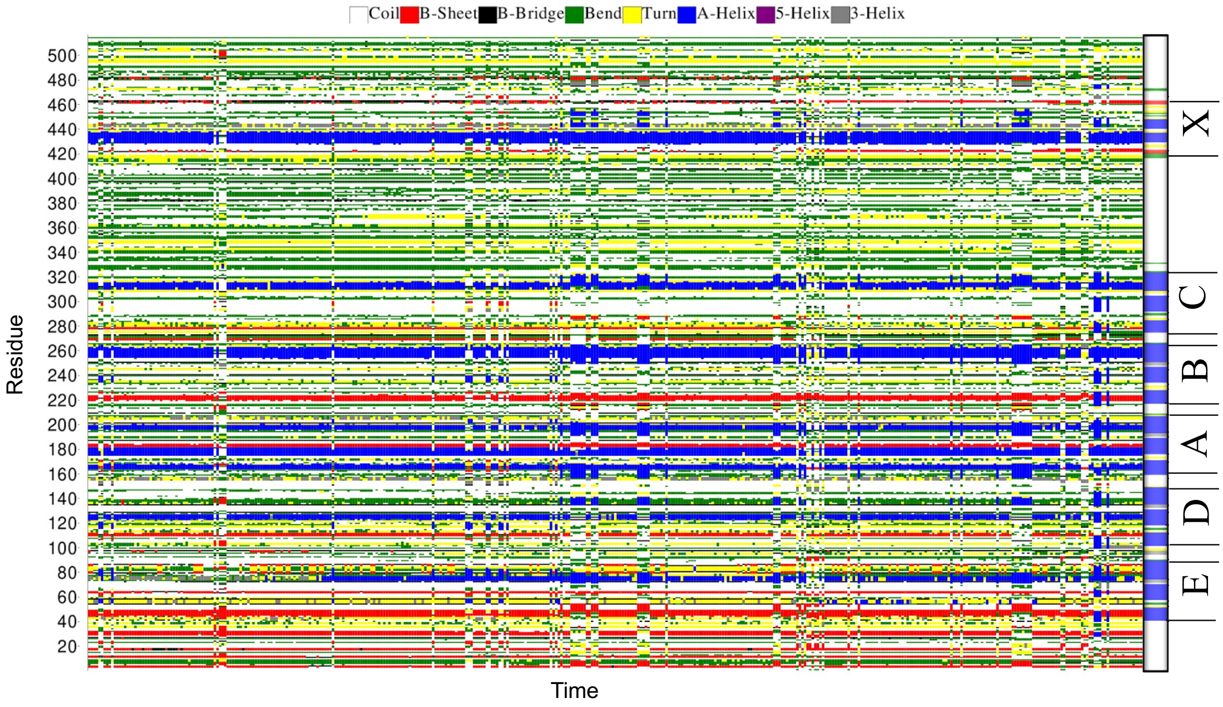


**Extended Data Fig. 10** | Color-coded DSSP analysis (not reweighted) as a function of simulation time for the subtrajectory $0.25 < Q < 0.35, 2.4 < R_g < 2.9$ nm of the main simulation. The rightmost bar displays the DSSP analysis of the AlphaFold-predicted structure, with domains E, D, A, B, C, and domain X labeled for reference.

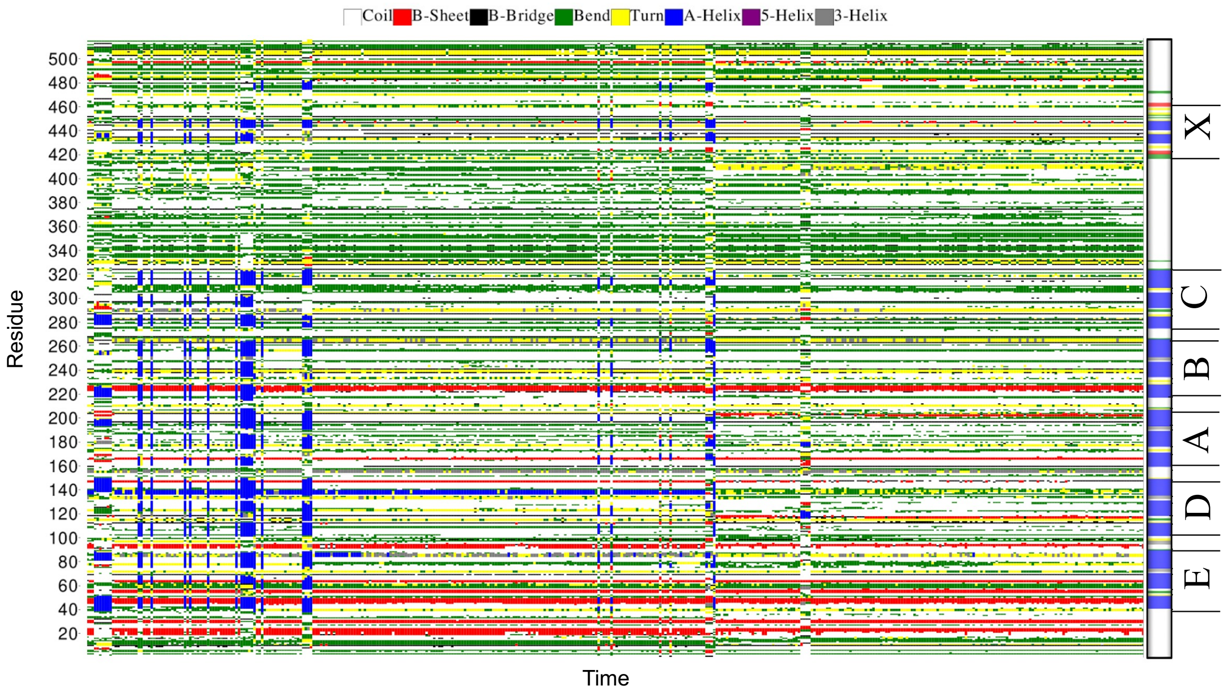


**Extended Data Fig. 11** | Color-coded DSSP analysis (not reweighted) as a function of simulation time for the subtrajectory $Q < 0.15, 2.4 < R_g < 2.85$ nm of the main simulation. The rightmost bar displays the DSSP analysis of the AlphaFold-predicted structure, with domains E, D, A, B, C, and domain X labeled for reference.

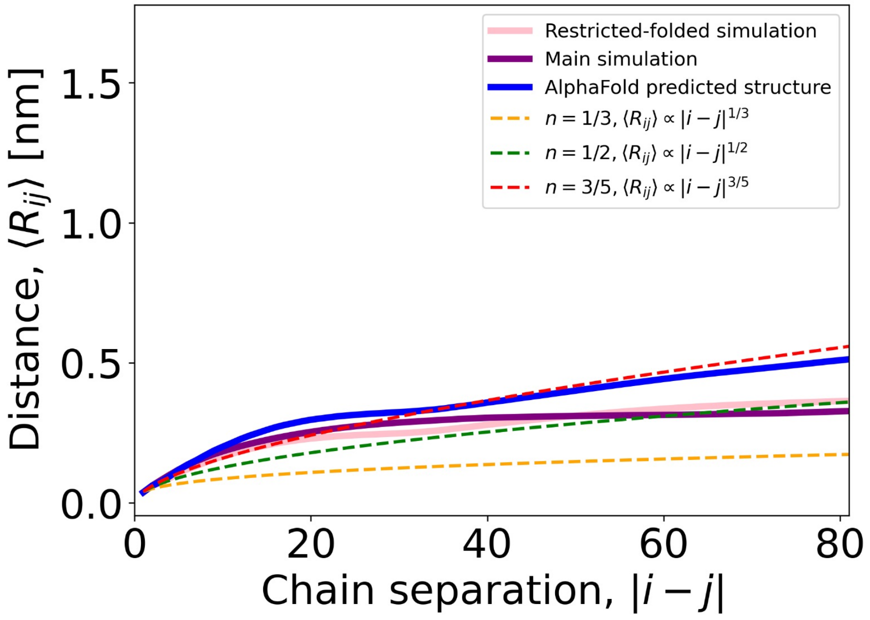


**Extended Data Fig. 12** | Reweighted ensemble-averaged interchain distance ($< R_{ij} >$) scalings for short chain separations (up to 80 residues) of full-length compared to three different Flory scaling exponents (dashed lines).

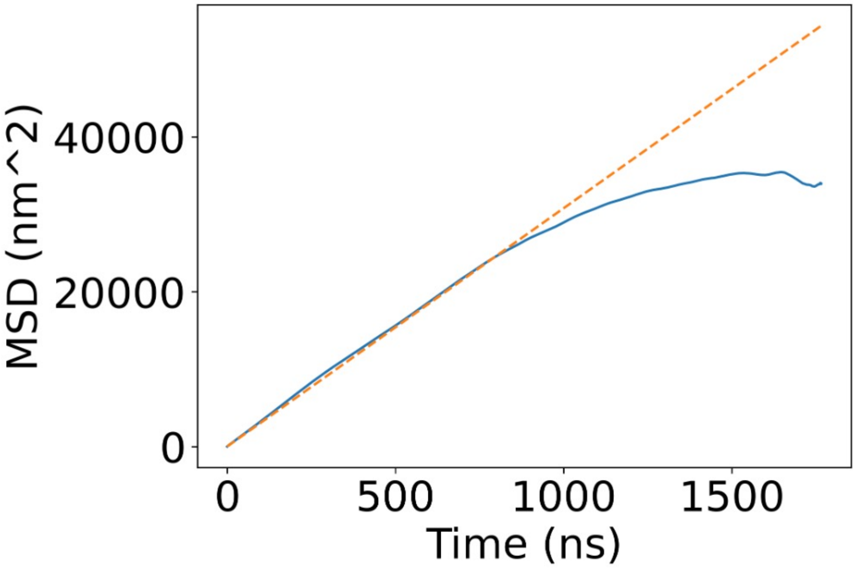


**Extended Data Fig. 13** | Mean square displacement (MSD) over time for the main simulation. The dotted orange line shows the best linear fit for the diffusivity.
